# Precision-targeted metabolomics method characterizes differential metabolomes in multiple-biological matrixes and cell-mitochondria

**DOI:** 10.1101/642496

**Authors:** Xialin Luo, Rui Guo, Jingjing Liu, Aihua Zhang, Xijun Wang, Haitao Lu

**Affiliations:** Key Laboratory of Systems Biomedicine (Ministry of Education), Shanghai Center for Systems Biomedicine, Shanghai Jiao Tong University, Shanghai 200240, China; Laboratory for Functional Metabolomics Science, Shanghai Jiao Tong University, Shanghai 200240, China; National Chinmedomics Research Center, Sino-America Chinmedomics Technology Collaboration Center, National TCM Key Laboratory of Serum Pharmacochemistry, Heilongjiang University of Chinese Medicine, Heping Road 24, Harbin, 150040, China

**Keywords:** precision-targeted metabolomics, functional metabolomes, multiple biological-matrixes, mitochondria metabolism, biomarker discovery

## Abstract

This new method has the capacity to dynamically analyse the metabolome of interest in diverse biological matrixes by offering coverage of rat urine, plasma, liver, brain, intestine, stomach, heart, spleen, lung, faeces, fresh plant tissues, cells and microbes. In addition, this new method enables specific and efficient analysis of microdontia metabolomes, non-microdontia and whole cell metabolomes, as well as can engage in absolute determination of 84 key clinical-wide metabolites in different biological matrixes, to enable the complementary support of clinical diagnosis and classification of diseases. To demonstrate the applicable capacity of this new method, multiple-matrixes differential metabolomes were firstly characterized using this new method to coordinate metabolic modifications underlie hepatitis induced by carbon tetrachloride (CCL_4_) in rats, such finding provides novel insight into the pathogenesis and therapeutics of hepatitis in clinic. Altogether, we are fully confident that this new metabolomics method will be widely welcomed by scientists in different niches to solve their key questions accordingly.

---

Metabolism is characterized by a series of essential life-sustaining processes in all organisms through which living cells acquire necessary nutrients and energy, which enables cells to grow, differentiate and function. By characterizing the altered metabolism underlying diverse biochemical events and/or processes, we could solve key problems in different niches, such as biomedicine, bioengineering, agriculture and the environment^1–3^. In the late 1990s, a systems biology-driven metabolomics method was first proposed to provide a comprehensive approach to precisely investigate metabolism^4,5^ via global and quantitative analyses of endogenous metabolites in biological systems that require data acquired through high-resolution analytical technologies^6^.

In terms of untargeted metabolomics always fails to precisely identify differential metabolites due to the undeveloped mass spectrometry database, and the limitations in the availability reference compounds ^7,8^. Targeted metabolomics is being prevalent and welcomed by the scientific community, which allows us to further explore the biochemical functions and mechanisms of dysregulated metabolism by engaging in functional experiments with the available reference compounds ^9,10^. Therefore, if we can establish a targeted metabolomics method with wide-coverage to important metabolites that is supposed to promote innovations in life sciences ^11,12, 13^.

According to our previous effort^13^, in this case, we selected 200+ important metabolites of interest to establish high-performance targeted metabolomics method **(Figure 1)**, which has a broad coverage to different metabolic pathways underlying different biological processes/events^13–16^. This newly developed targeted metabolomics method was seriously defined as a high-performance method by reducing all the unstable and low-resolution metabolite signals, enabling to collect high-quality metabolomics data accordingly ^17–18^ **(Figure 3).**

**Figure 1.**
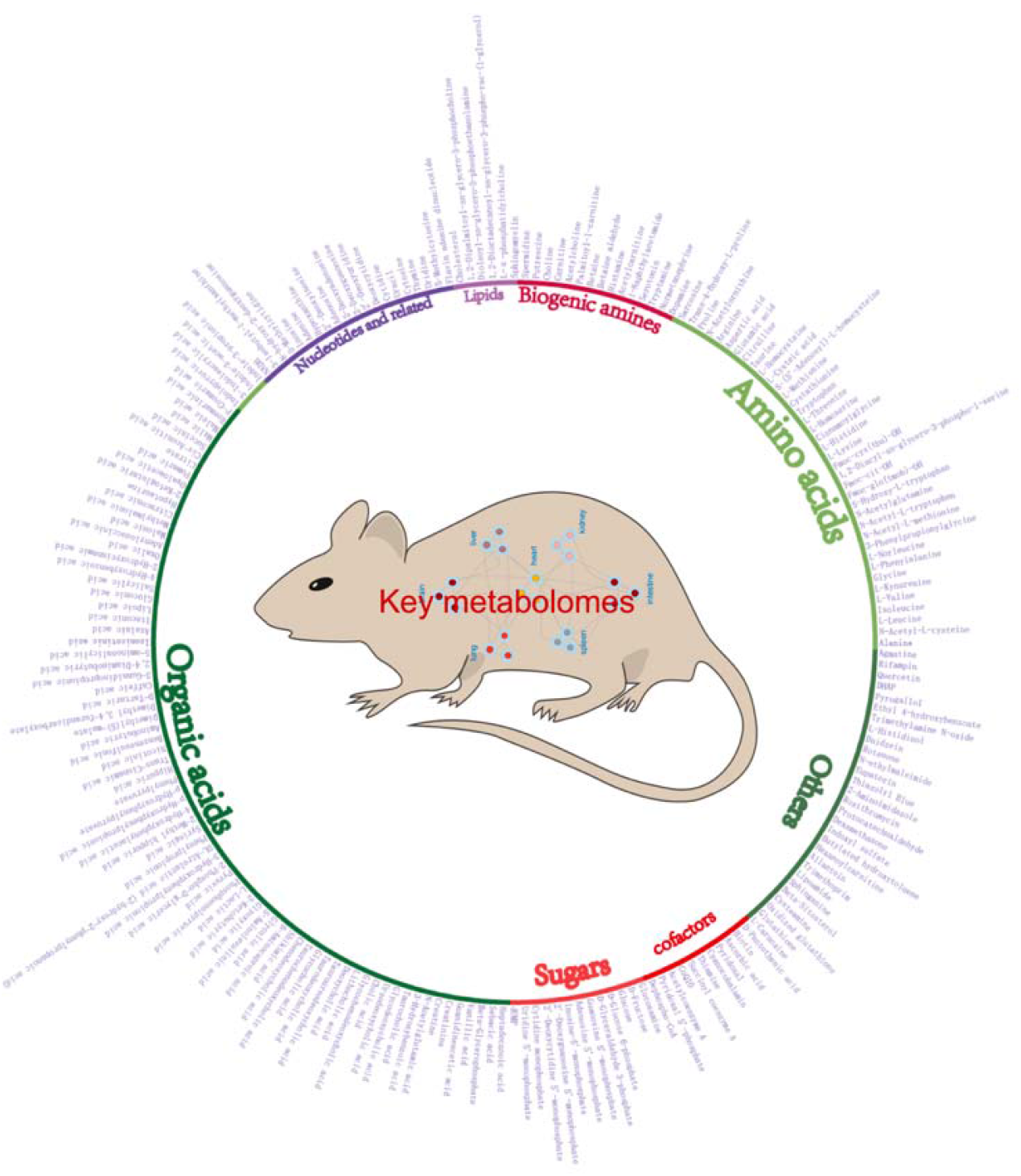
240 key metabolites with importantly biochemical functions are captured by the proposed targeted metabolomics method.

**Figure 2.**
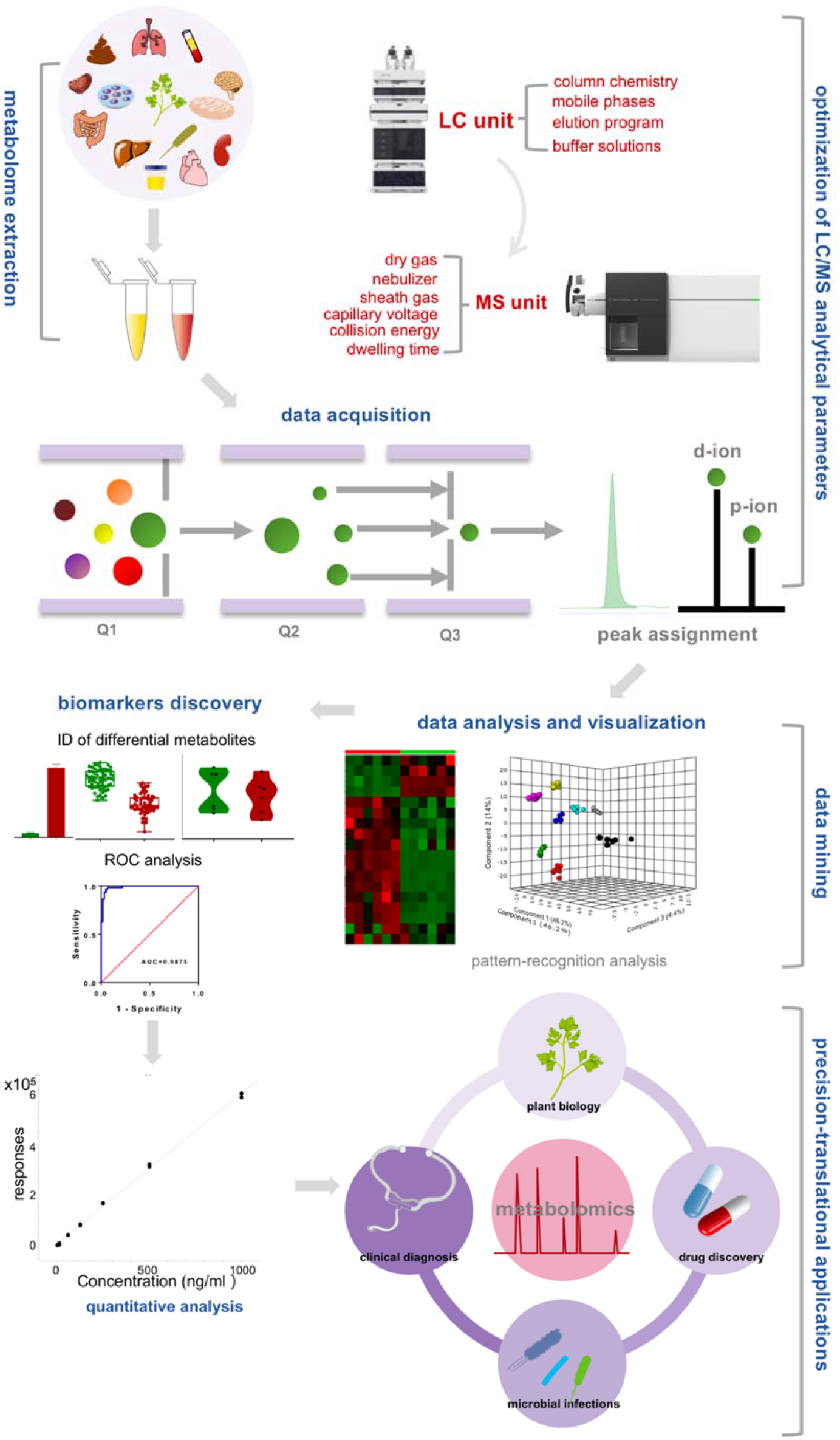
schematic illustration of the developed targeted-metabolomics method from the optimization of analytical parameters, high-quality data-acquisition, data analysis/visualization, absolute determination of key metabolites to the capacity of enabling translational applications in different niches of life sciences.

**Figure 3.**
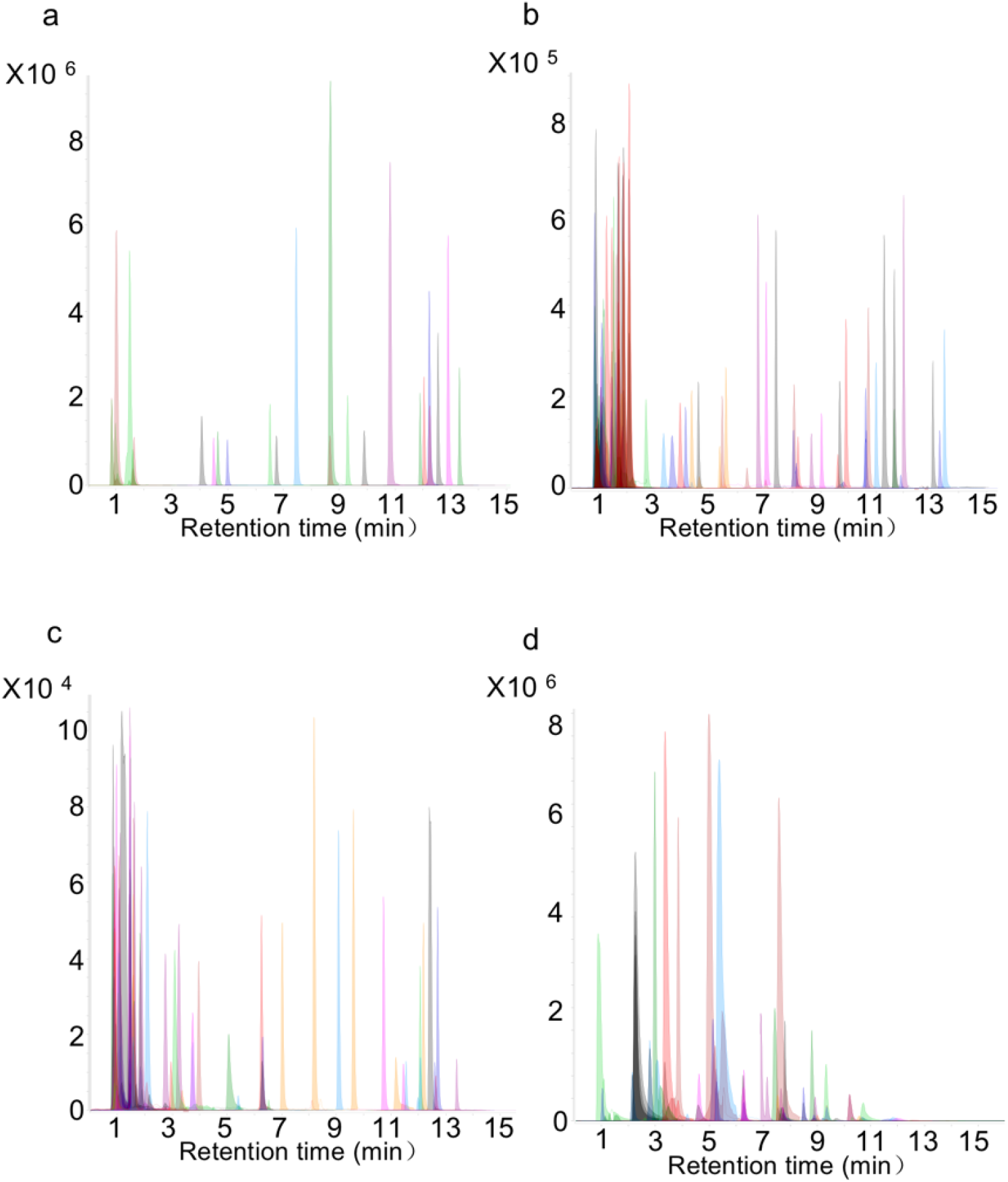
Typical reference-compound EIC profiles for 240 targeted metabolites collected by both by reverse-phase T3 column (A, B and C), and normal-phase HILIC column (D). **Figure 3** schematic illustration of the developed targeted-metabolomics method from the optimization of analytical parameters, high-quality data-acquisition, data analysis/visualization, absolute determination of key metabolites to the capacity of enabling translational applications in different niches of life sciences.

To tailor the translational applicability of this method for use in the full-spectrum of life sciences, we sought to analyse the metabolomes of interest present in a range of diverse biological matrixes **(Figure 4).** It is exciting to illustrate the collective data that this protocol has been able to obtain efficiently for targeted characterization of metabolomes from rat urine, plasma, liver, brain, intestine, stomach, heart, spleen, lung and faeces; fresh plant tissues; cells; and microbes **(Figure 5A and C).**

**Figure 4.**
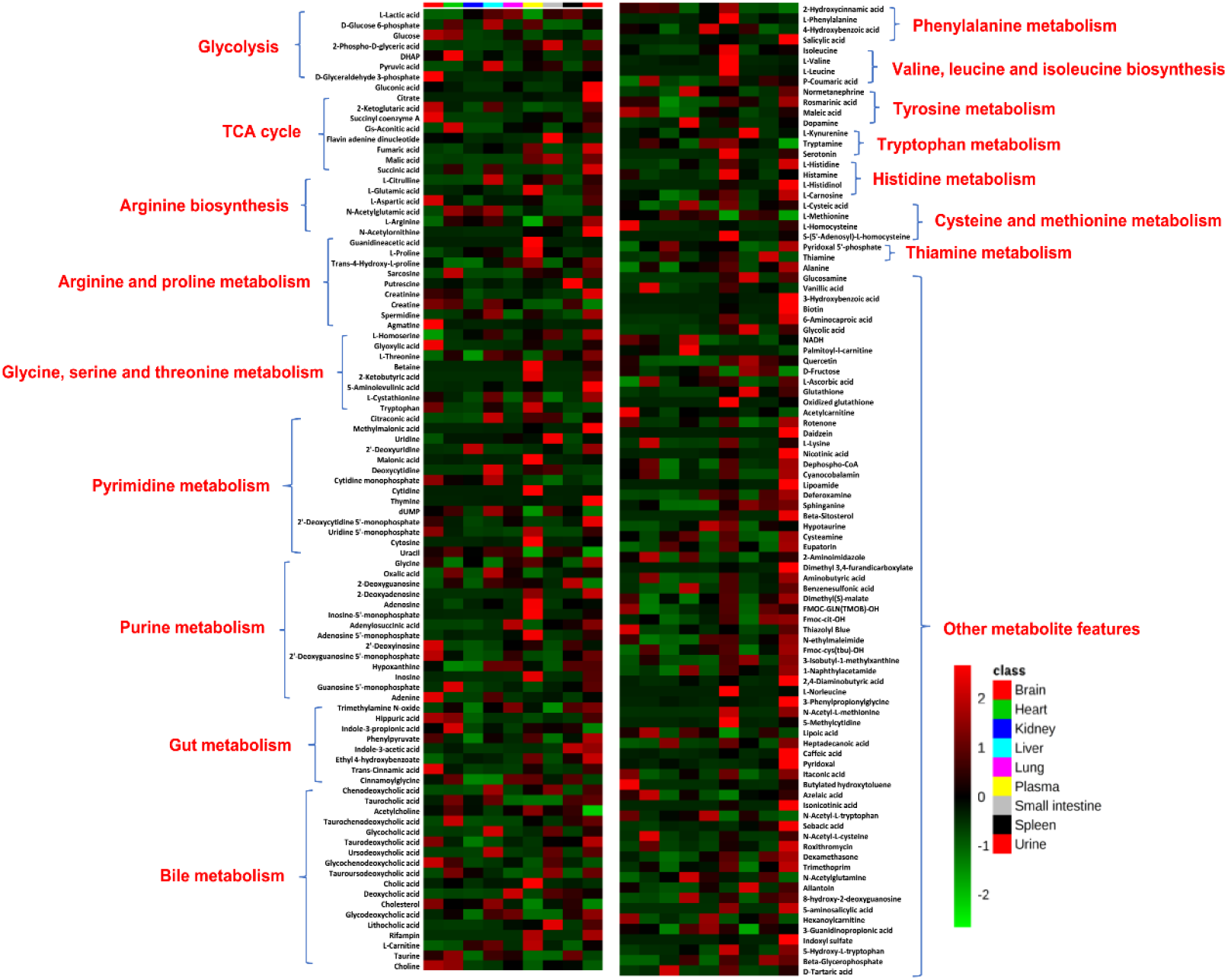
Heatmap overview of differential metabolomes with selected tissues

**Figure 5.**
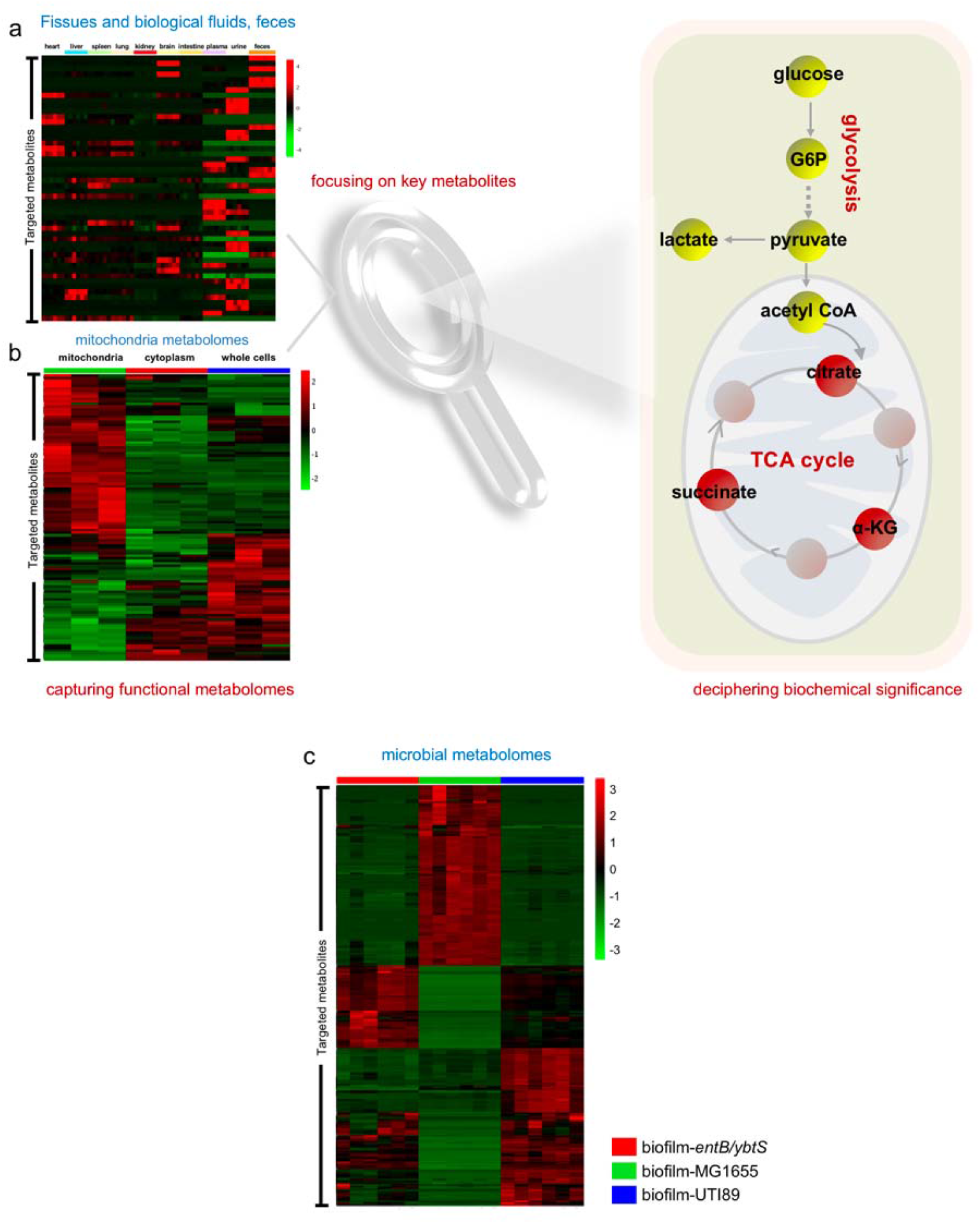
The targeted metabolomics method can be broadly used for precision-assay of differential metabolomes in differently biological-matrixes, including tissues, biological fluids and feces (A); mitochondria, cytoplasm and whole cells (B); and microbial organisms (C).

Importantly, in terms of microdontia, it is quite important to understand the related cell metabolism, because metabolic dysregulation is implicated in numerous important biological processes and events, such as cancers, diabetes, and immunological deficiency, and the whole cell-metabolome assay often cannot be used for precise characterization of the functional features of dysregulated microdontia related to different biological events. To improve this assay, our new protocol was developed to enable the simultaneous analysis of microdontia metabolomes, non-microdontia metabolomes, and whole-cell metabolomes **(Figure 5B)**. When making a comparison among cancer cell lines, microdontia metabolomes are much better than non-microdontia metabolomes or whole-cell metabolomes for investigating metabolic reprogramming during cancer progression. Finally, for improved clinical diagnosis and classification of targeted diseases, we also determined calibration curves for precise quantification of metabolite biomarkers, and 84 key metabolites were eventually and absolutely quantified in both plasma and urine samples, as well as other tissue matrixes **(Tables 2 and 3).** Collectively, the rapid determination of these 84 key metabolites with our targeted metabolomics protocol has great potential for use in complementing the clinical diagnosis of complex diseases **(Figure 6)**. To demonstrate the applicable capacity of this new method, multiple-matrixes differential metabolomes were firstly characterized using this new method to coordinate metabolic modifications underlie hepatitis induced by carbon tetrachloride (CCL_4_) in rats. Such findings verify the robustness and applicable potential of this new targeted metabolomics method, as well as provide novel insight into the pathogenesis and therapeutics of hepatitis in clinic from multisensorial metabolic perspective **(Figure 7).**

**Figure 6.**
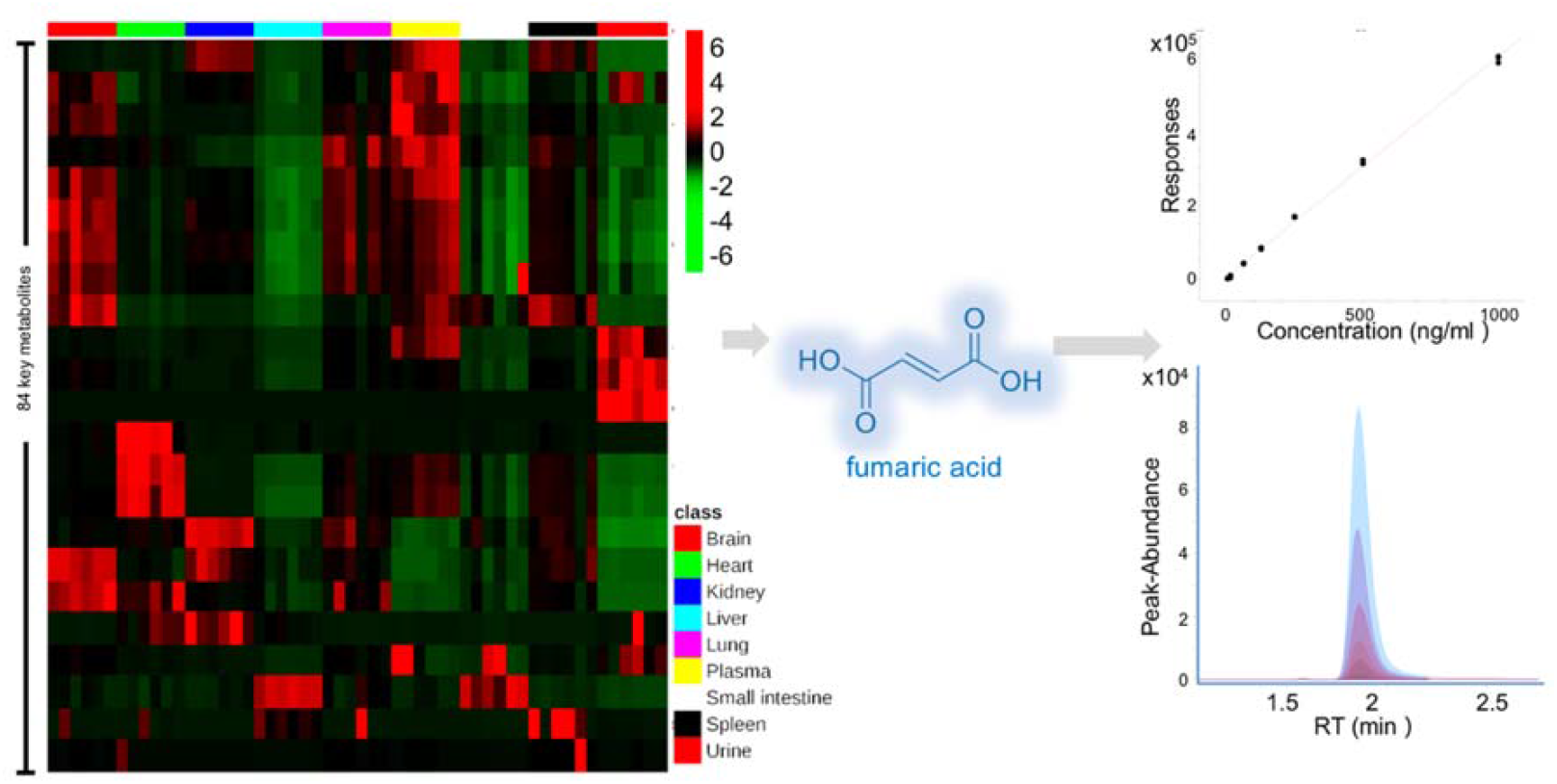
84 clinical-wide key metabolites can be absolutely determined from different biological matrixes with our new targeted-metabolomics method, particularly urine and plasm. Such effort can complementarily support clinical diagnosis of numerous diseases.

**Figure 7.**
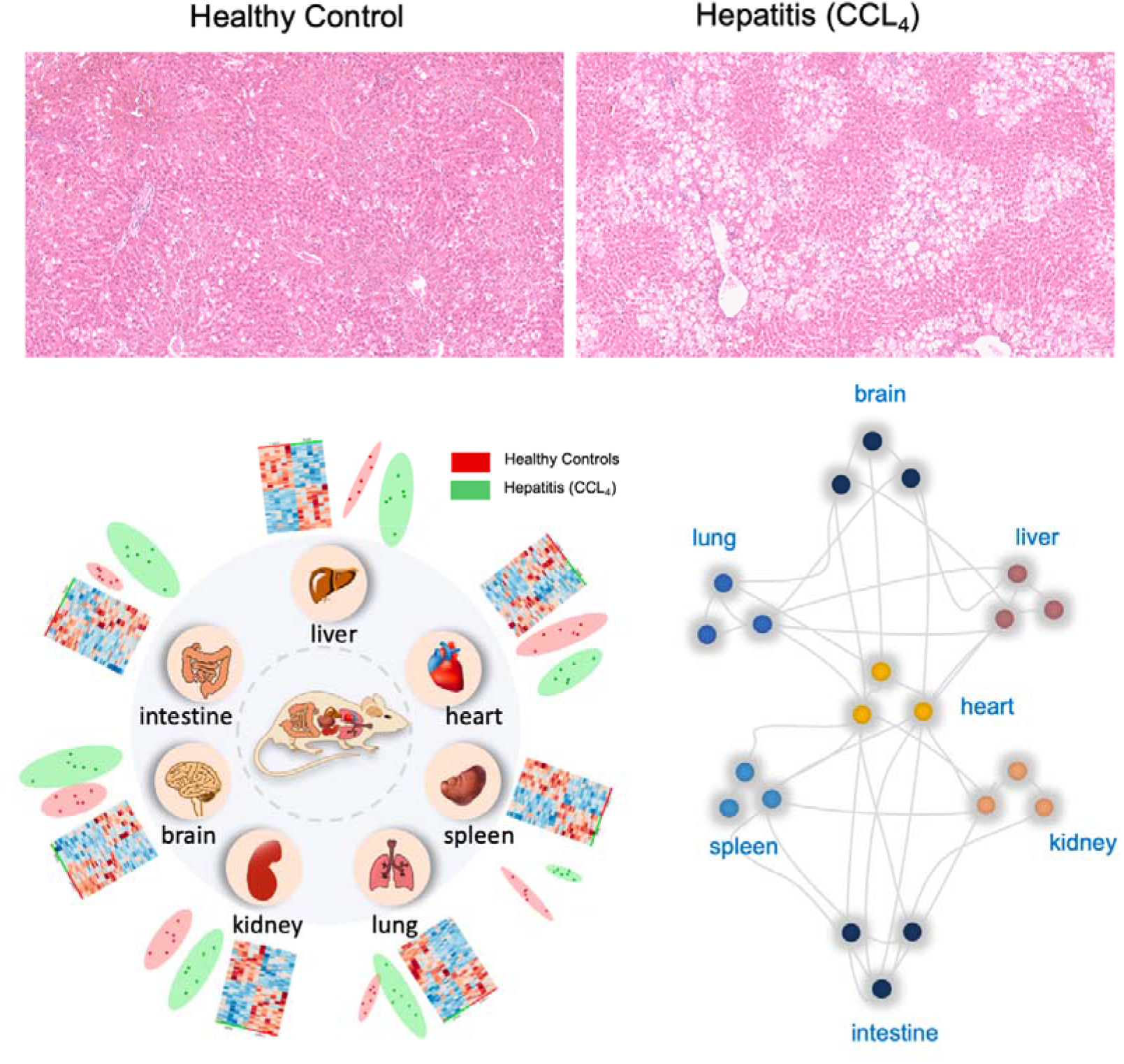
this method is used to characterize metabolic pattern throughout multiple tissues of hepatitis induced by CCL_4_ in rats. The data completely reveals high-resolution metabolic modifications of hepatitis that are closely associated with different tissues besides liver. Such findings verify the robustness and applicable potential of this new targeted metabolomics method, as well as provide novel insights into unpredictable metabolic features of hepatitis to enable future pathogenesis annotation and therapeutic discovery.

Taking all these advantages together, this novel protocol for targeted metabolome assays has substantial coverage of 240 key metabolites in dozens of important metabolic pathways. This method can enable the use of metabolome assays with different biological matrixes and offers a new horizon for analyses of microdontia metabolomes, non-microdontia metabolomes, and whole-cell metabolomes.

In short, this new method is far better than previous protocols of targeted metabolome assays, particularly in terms of matrix adaptation, capacity for absolute quantification, and new and future applications for the analysis of microdontia metabolomes and non-microdontia metabolomes. We are fully confident that this protocol will be widely welcomed by scientists studying different biological niches to solve their key questions accordingly **(Figure 8).**

**Figure 8.**
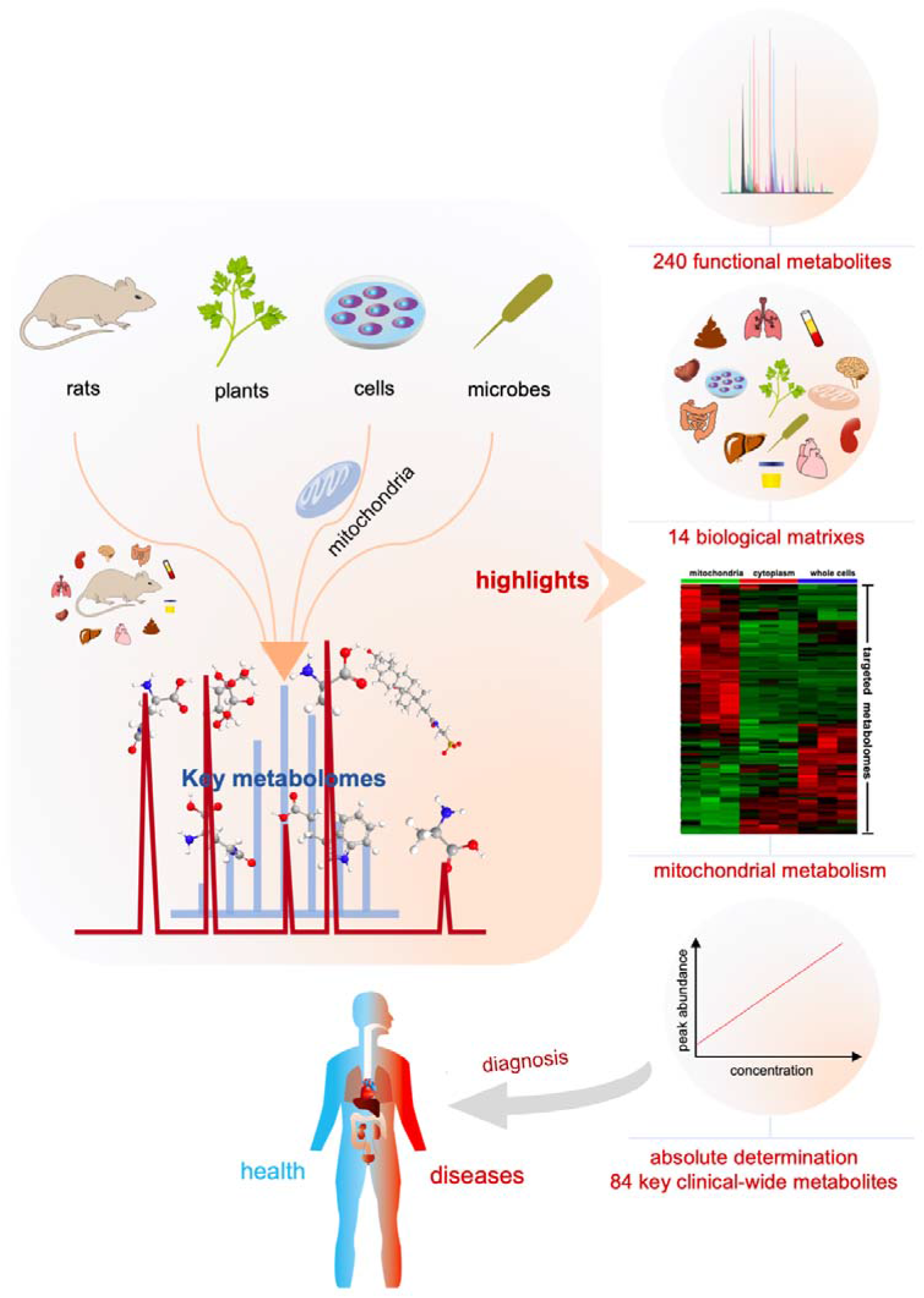
Unique features of this new targeted metabolomics method are highlighted as 1) using an RPLC column for a hydrophilic metabolome assay and a HILIC column for a hydrophobic metabolome assay. 2) with our new targeted metabolomics protocol, we substantially increased the coverage to 240 metabolites of interest with largely variable polarity and improved detection capacity. 3) this new protocol has the capacity to dynamically analyse the metabolome of interest in diverse biological matrixes by offering coverage of rat urine, plasma, liver, brain, intestine, stomach, heart, spleen, lung and faeces; fresh plant tissues; cells; and microbes. 4) this protocol enables specific and efficient analysis of microdontia metabolomes, non-microdontia and whole cell metabolomes. 5) absolute determination of 84 key metabolites in different biological matrixes enables the complementary support of clinical diagnosis and classification of diseases.

## Materials and Methods

### Chemicals and reagents

All of reference compounds for targeted metabolomes of interest (**Table 1**) were purchased from Sigma-Aldrich Corp (Saint Louis, USA), and 1 mg/mL stock solution was prepared by dissolving them into water or methanol, respectively. Methanol, acetonitrile and formic acid (HPLC-grade) were purchased from Thermo Fisher Scientific (Shanghai, China). Ammonium acetate (HPLC grade) was purchased from Merck KGaA (Darmstadt, Germany). LB broth (Luria−Bertani) and LB agar were purchased from Becton Dickinson (Sparks, USA). Magnesium sulfate, manganese chloride, casamino acid, yeast extract, of USP grade, were purchased from Sigma-Aldrich Corp (Saint Louis, USA). The distilled water was purchased from A.S. Watson Group (Guangzhou, China). All other reagents were of ACS grade.

### UPLC/TQ MS system

An UPLC system (1290 Infinity series, Agilent Technologies) coupled to a triple quadrupole mass spectrometer (Agilent 6495, Agilent Technologies) was used for precision-targeted metabolome assay in a dynamic MRM mode (DMRM). The optimized dynamic MRM parameters for the targeted metabolomes were listed in **Table 1**. The key parameters for MS unit are briefly summarized as, the ion source was Agilent Jet Stream ESI and the parameters were set as follows: dry gas temperature, 250 °C; dry gas flow, 16 L/min; nebulizer pressure, 20 psi; sheath gas heater, 350 °C; sheath gas flow, 12 L/min; capillary voltage, capillary voltage, 4000 V for positive mode and 3500 V for negative mode. RPLC separation was performed with an ACQUITY UPLC HSS T3 (2.1×100 mm, 1.8 μm) column; the mobile phase A was water with 0.1% formic acid (v/v) and mobile phase B was acetonitrile with 0.1% formic acid (v/v) with an optimized gradient-elution program: 0-2 min, 98% A; 2-10 min, 98%-65% A; 10-12 min, 65%-20% A; 12-14 min, 20%-2% A; 14-30 min, 2% A. HILIC separation was carried out with an ACQUITY UPLC BEH Amide column (2.1 mm i.d×100 mm, 1.7 μm; Waters), the mobile phase A is changed to water with 0.1% (v/v) formic acid and 10mM ammonium acetate, while mobile phase B is acetonitrile with 0.1% formic acid (v/v); the relevant gradient-elution program was optimized as: 0-4 min, 5% −12% A; 4-15min, 12%-50%; 15-25 min, 50% A. The flow rate for both RPLC and HILIC separation systems was set at 0.3 mL/min and the column temperature was maintained at 40 °C. All the samples were placed in 10 °C with a 5 μL-volume injection.

### Rat sampling and collection

Male SD rats (age: 6 weeks, body weight: 160-200 g) were supplied by GLP Center of Heilongjiang University of Chinese Medicine (Harbin, China). In treated group, 6 rats were orally intraperitoneal injected with 20% CCl_4_ olive oil solution according to the rat weight at 5 mL/kg, to induce liver injury, another 6 rats were treated with olive oil solution as healthy controls. Urinary samples were collected with metabolic-cage and centrifuged under 13000 rpm at 4 °C for 15 min. The rats were sacrificed by exsanguination after anesthetized by intraperitoneal injection of 3% pentobarbital sodium according to the rat weight at 2 mL/kg, and the plasma samples were collected and centrifuged under 3500 rpm at 4 °C for 15 min. In addition, the tissues, such as brain, heart, liver, spleen, kidney, small intestine, and lung samples were collected and cleaned in normal saline, then rapidly frozen in liquid nitrogen. Finally, tissues, urine and plasma samples were all stored at −80 °C until use. All animal care and experimental procedures were performed in compliance with the rules on the care and use of animals of the Ethical Committee of Heilongjiang University of Chinese Medicine. All efforts were made to ameliorate the suffering of the animals.

### Bacterial strains and cultivation

The UTI89 and MG1655 strains were firstly cultured in LB-agar plate for 12 h, then one colony was collected to incubated with LB broth medium for 4 h, diluted 1000-fold into CFA medium and further cultivated for 72 h at 30°C to trigger mature biofilm formation. CFA medium is consisted of 1% Casamino acids, 0.15% yeast extract, 0.005% magnesium sulfate and 0.0005% manganese chloride.

### Isolation of mitochondria from cells

Cell line Panc-1 was purchased from the American Type Culture Collection (ATCC), the cells are cultured in DMEM medium supplied with 10% fetal bovine serum and 1% Penicillin-Streptomycin, maintained at 37°C in a humidified atmosphere containing 5% CO_2_. In order to obtain a significant quantity of mitochondria, an appropriate number at about 3 million cells were seeded in a 150 mm dish and grown to approximately 2 × 10^7^ cells. The isolated mitochondria and cytosolic fraction were achieved by commercially available mitochondria isolation kit (Beyotime, China). The isolation was performed according to a modified manufacturer’s protocol. Briefly, cells were washed with ice cold PBS, followed by adding 1 mL of the prepared 1×Extraction Buffer A, then incubated on ice for 10 minutes. To further homogenize the cells on ice with a dounce homogenizer for 15 strokes. The extraction was centrifuged under 600 × g for 10 minutes at 4 °C, and carefully transfer the supernatant into a new PE tube, centrifuge under 11,000 × g for 10 minutes at 4 °C. At last, the supernatant was collected as cytosolic fraction and the pellets were mitochondria. All the isolation process should be performed on ice.

### Metabolome extraction

#### Body fluids

Referring to our previous protocol^13^, we made a revised version as below, briefly, the urine samples could be directly analyzed with LC-TQ MS after a simple preparation as urinary samples were centrifuged under 20,000 g at 4 °C for 10 min and filtered by 0.22 μm Millipore filter. 100 uL of plasma samples were mixed with 4 volume of iced-cold acetonitrile to precipitate proteins, then centrifuged to 20,000 g for 10 min at 4 °C. Following the supernatant was transferred to a new 1.5-mL microcentrifuge tube and dried with N_2_, finally dissolved in 100 μl of water and centrifuged again before analyzed by LC-TQ MS.

#### Tissues

Brain, heart, liver, spleen, kidney, small intestine and lung share the same protocol for metabolome extraction. About 100-130 mg tissue samples were weighed and placed into a 2-mL screw-cap plastic microvials containing about 1 g glass beads (1.0 mm i.d.). Then tissues were completely homogenized at a 2-min run for 3 times (after one run we need to put the sample on ice for cooling) in 1.2 ml of 80% ice-cold methanol by Mini-Beadbeater-16 (Biospec Products). After that, spun down at 20,000 g for 10 min and vortexed with 800 μl of iced-cold acetonitrile, centrifuged again to remove the protein precipitates. The supernatant was processed following the same procedure as plasma samples. Quality control (QC) samples by pooling aliquots of all samples were analyzed during the whole LC-TQ MS analysis.

#### Bacterial strains

Referring to the previous protocol^14^, we made a revised version. Firstly, biofilm samples were isolated from 50 mL of culture solution, to obtain the planktonic cells, the supernatants in the culture medium were centrifuged to 931 g at 4 °C for 15 min. After wash the biofilm with 2 mL PBS for 3 times, add 2 mL of 80% ice-cold methanol, homogenize by homogenizer for 2 minutes, 3 times. All procedures were performed on ice for cooling. Then, the homogenate was spun down at 13201 g to collect the supernatants. After adding 800 μL of ice-cold acetonitrile, centrifuged again, the fresh supernatants were filtered through 0.22μm filter membrane and stored in −80℃before lyophilization. The lyophilized samples were dissolved in 100 μL of water and centrifuged again before analyzed with LC-TQ MS.

#### Cells, cytoplasm and mitochondrial

metabolome extraction of the isolated mitochondrial was done by following the same procedure as that of tissue samples except for replacing the 1.0 mm i.d. of glass beads with 0.5 mm. The cytoplasm samples were mixed with 800 μL iced-cold acetonitrile to precipitate proteins, then following the steps of plasma-metabolome preparation. For whole cells, firstly, wash cells twice with PBS, following add 1.2 mL 80% ice cold aqueous methanol and all the cells were harvested by cell scraper on ice, and transferred to a 2-mL screw-cap plastic micro-vial, then homogenized, centrifuged; at last, adding iced-cold acetonitrile and following the routine procedure as the metabolome extraction of bacterial cells.

### Absolute determination of key clinical-wide metabolites

To assist clinical diagnosis of a variety of diseases with this new developed metabolomics method, we adopted a set of external standards by containing at least five-levels concentration to establish the calibration curve. 84 key clinical-wide metabolites can be absolutely determined by the developed precision-targeted metabolomics method **(Table S2)**. In this case, we attempt to verify the efficiency of this new developed method by selectively determining the absolute concentration of 27 key metabolites present in a diverse biological-matrixes **(Table S3)**.

### Data Processing and Statistical Analysis

The MS raw data generated from the biological samples was firstly processed by qualitative Analysis, with an in-house software of Agilent Corps, which integrated the signal-peaks and generated a three-dimension data-matrixes with peak-area of merabolite-metabolite ID-sample ID. After peak area for each metabolite was normalized to tissue weight of each sample, the processed data was uploaded onto free MetaboAnalyst online-softare (http://www.metaboanalyst.ca/MetaboAnalyst/) for partial least-squares discriminant analysis (PLS-DA) and heatmap vs hierarchical-cluster analysis (HCA).

## Acknowledgements

This work was supported by the National Key R&D Program of China (No. 2017YFC1308600 and 2017YFC1308605), the National Natural Science Foundation of China Grants (No. 81274175 and 31670031), the Startup Funding for Specialized Professorship Provided by Shanghai Jiao Tong University (No. WF220441502), and the Fundamental Research Funds for the Central Universities (grant no. 106112015CDJZR468808).

---

**Table S1.**
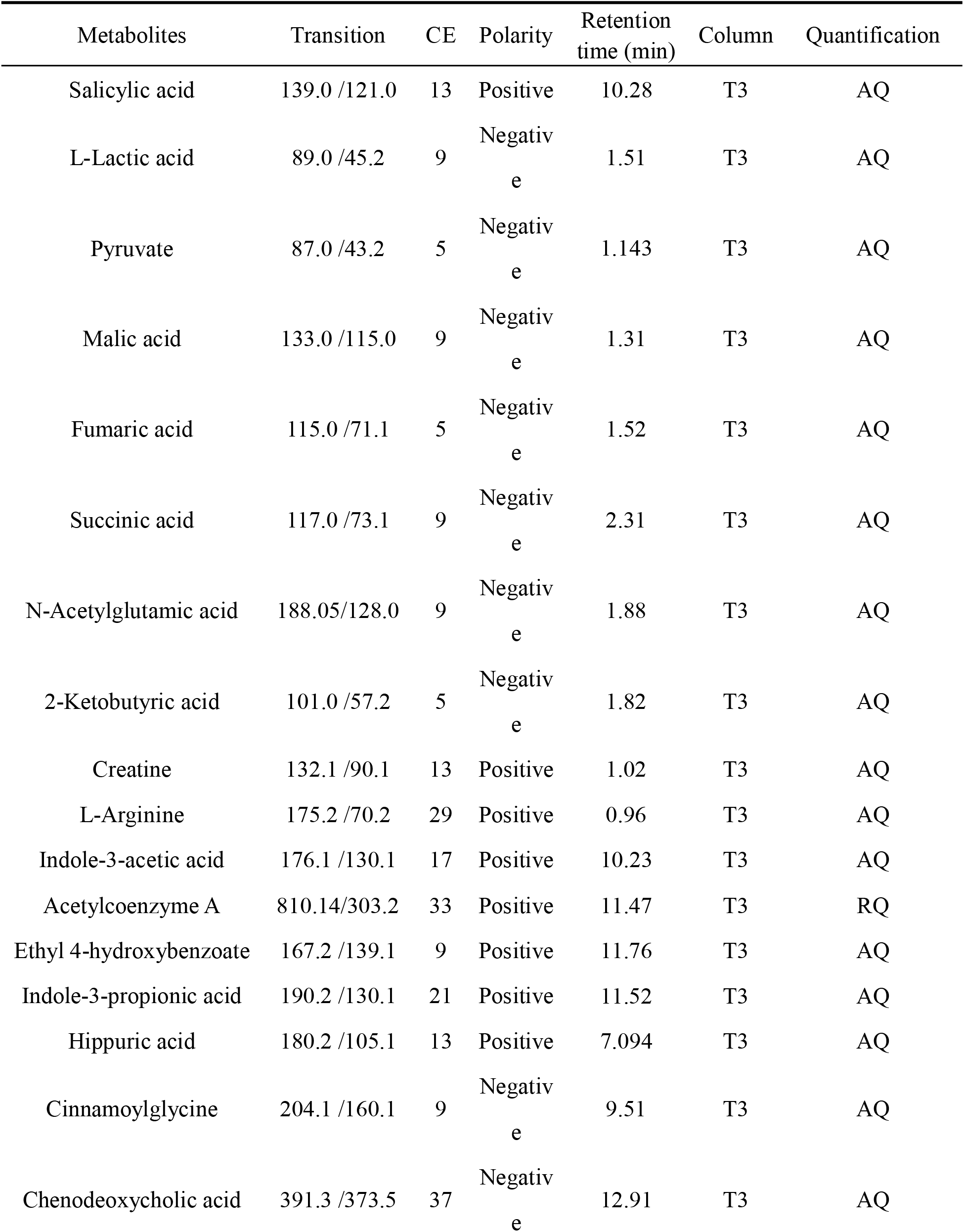
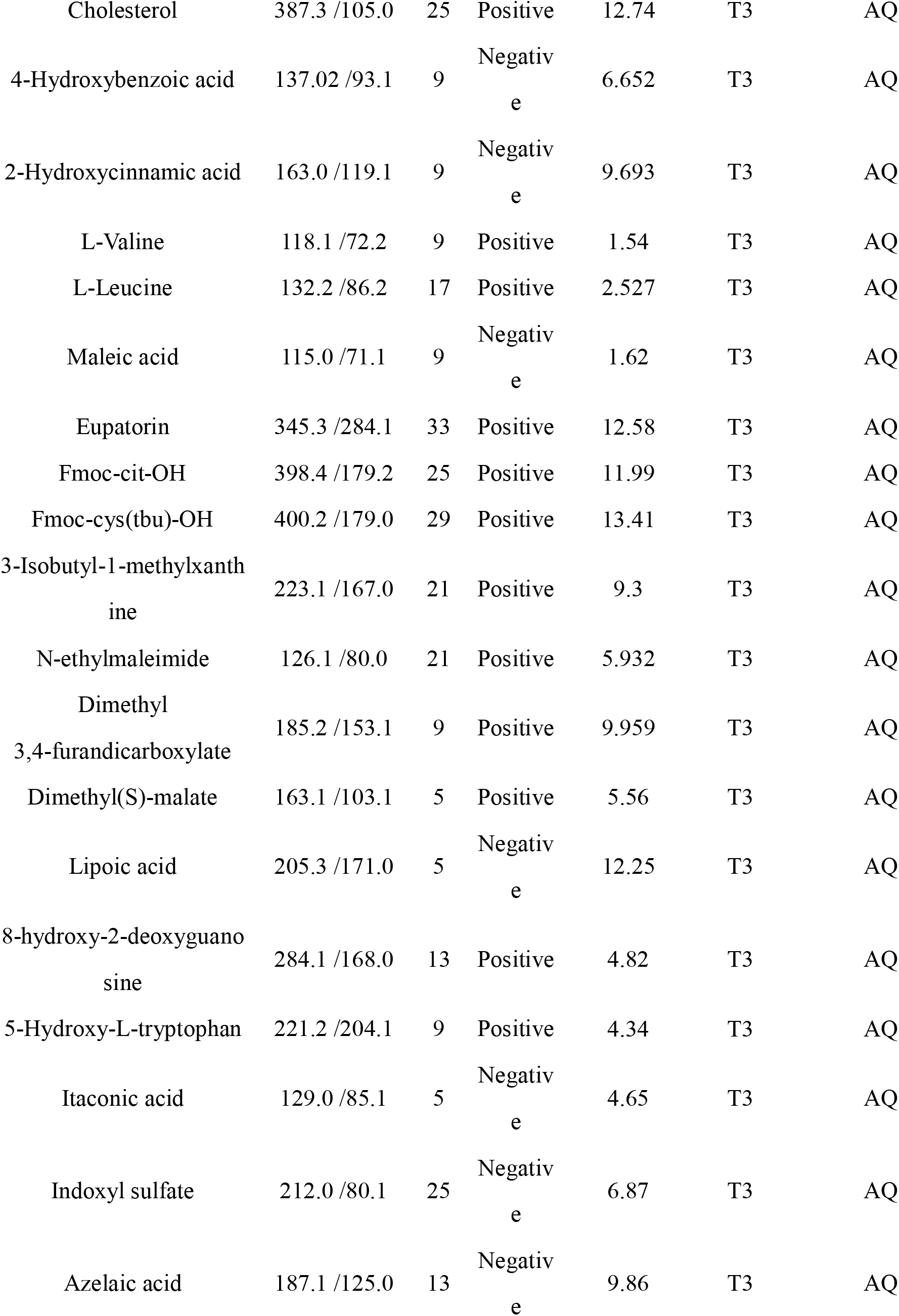
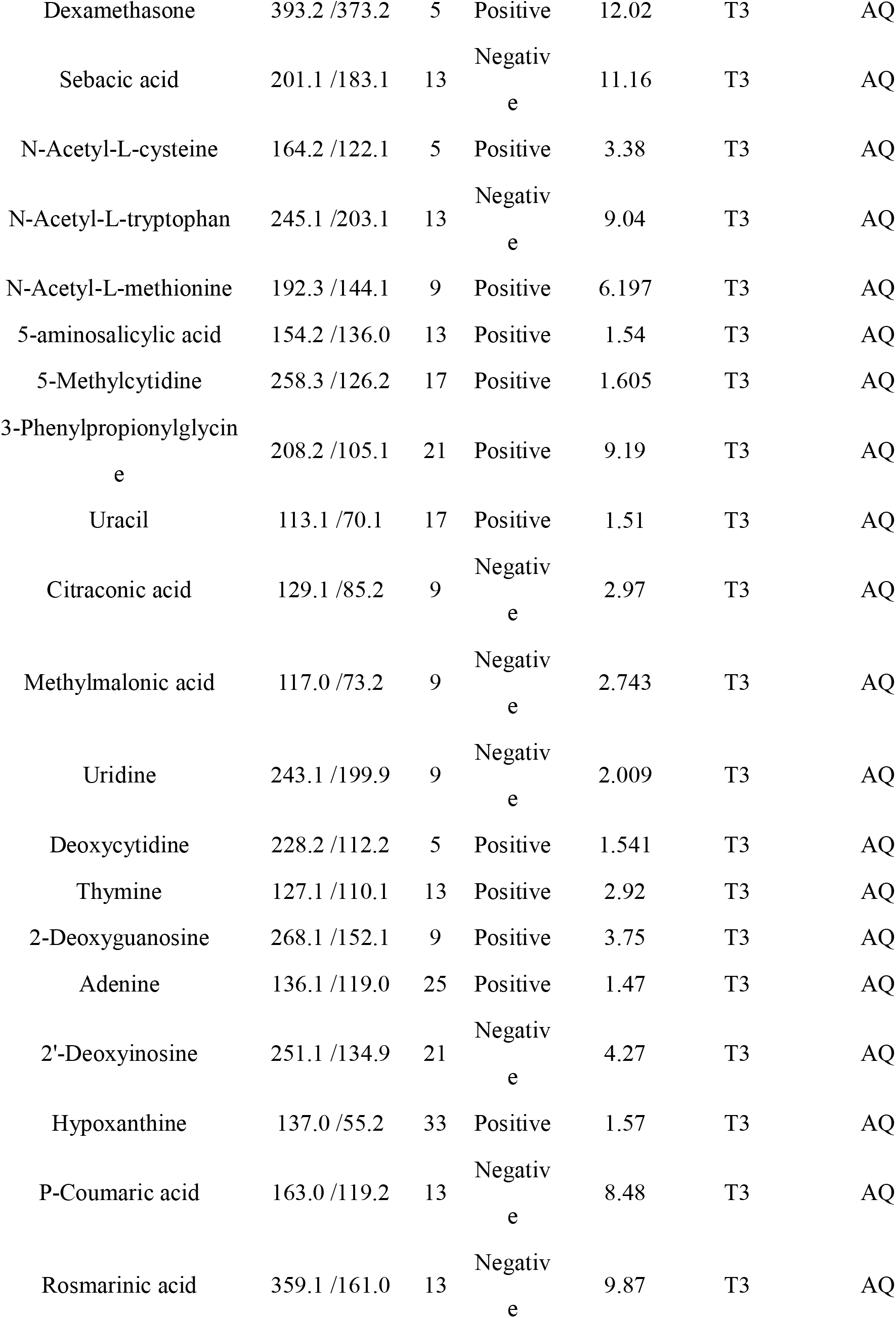
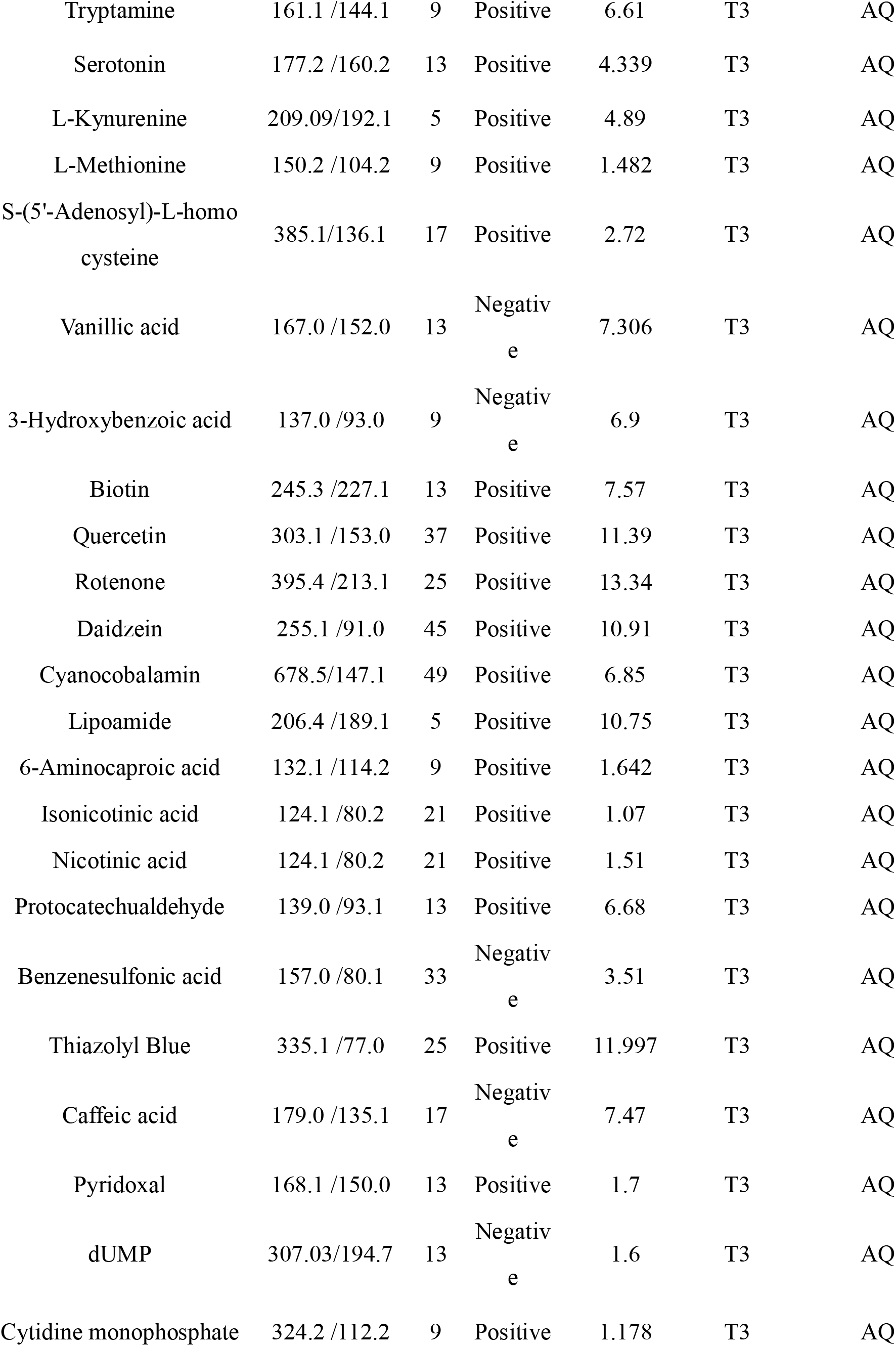
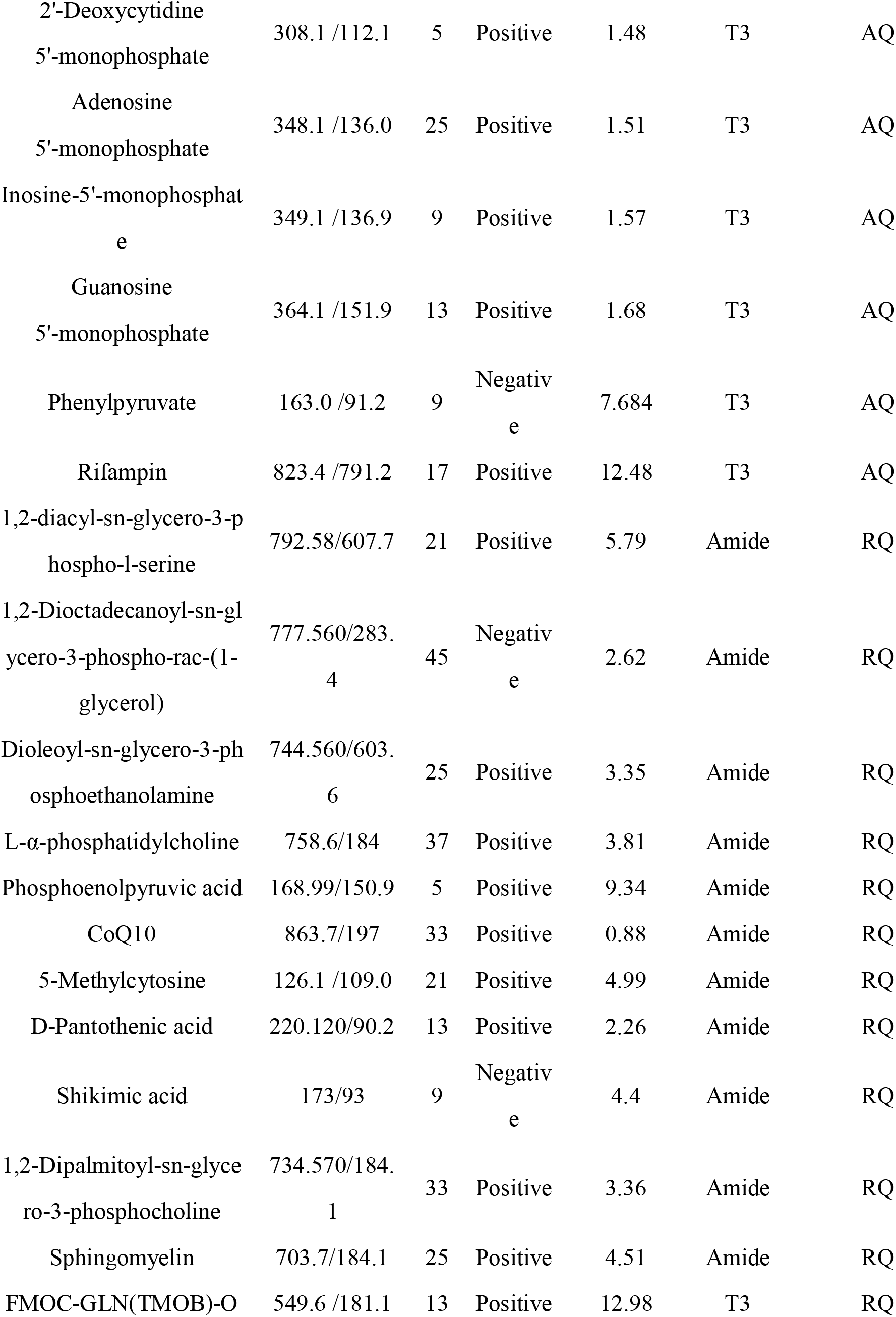
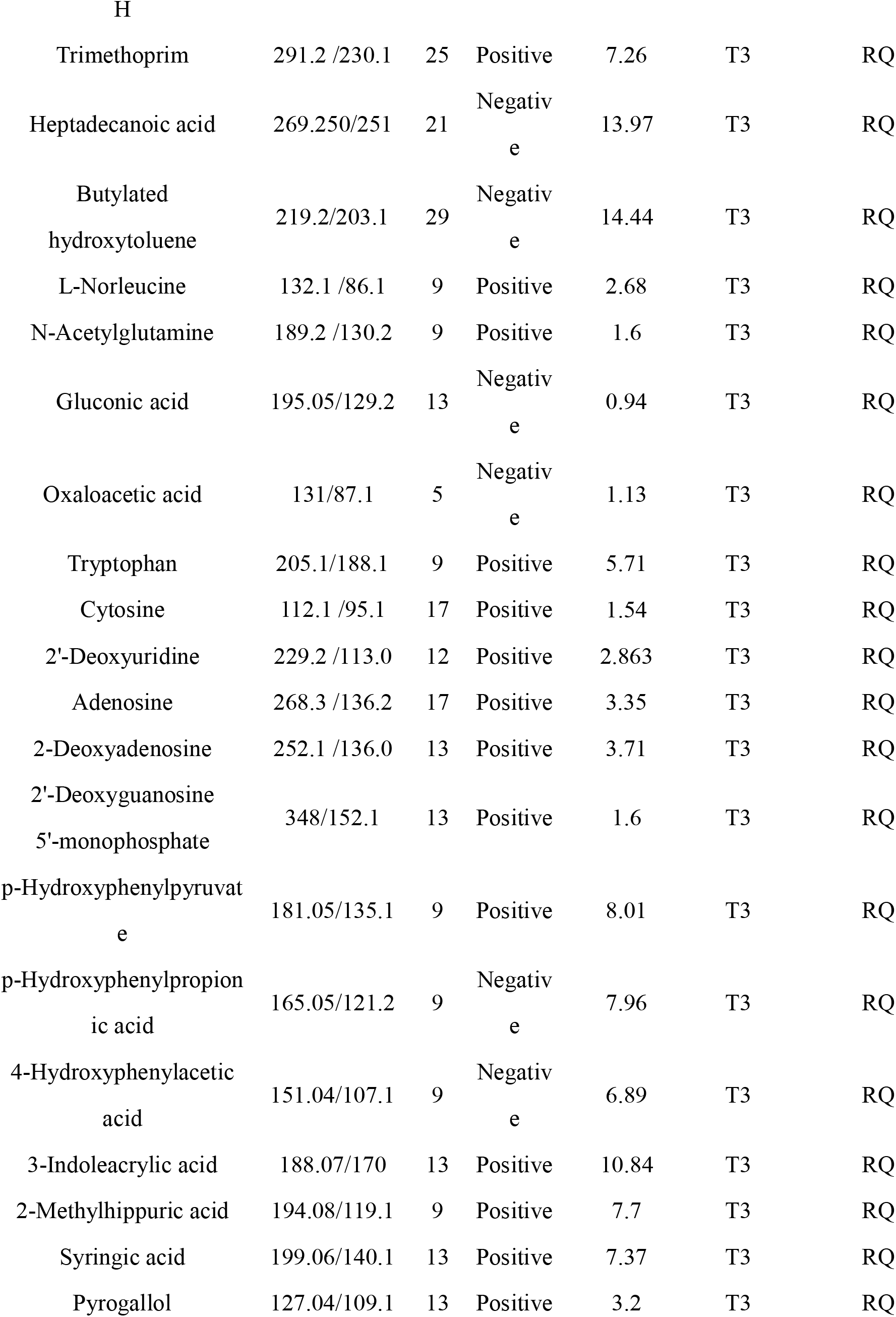
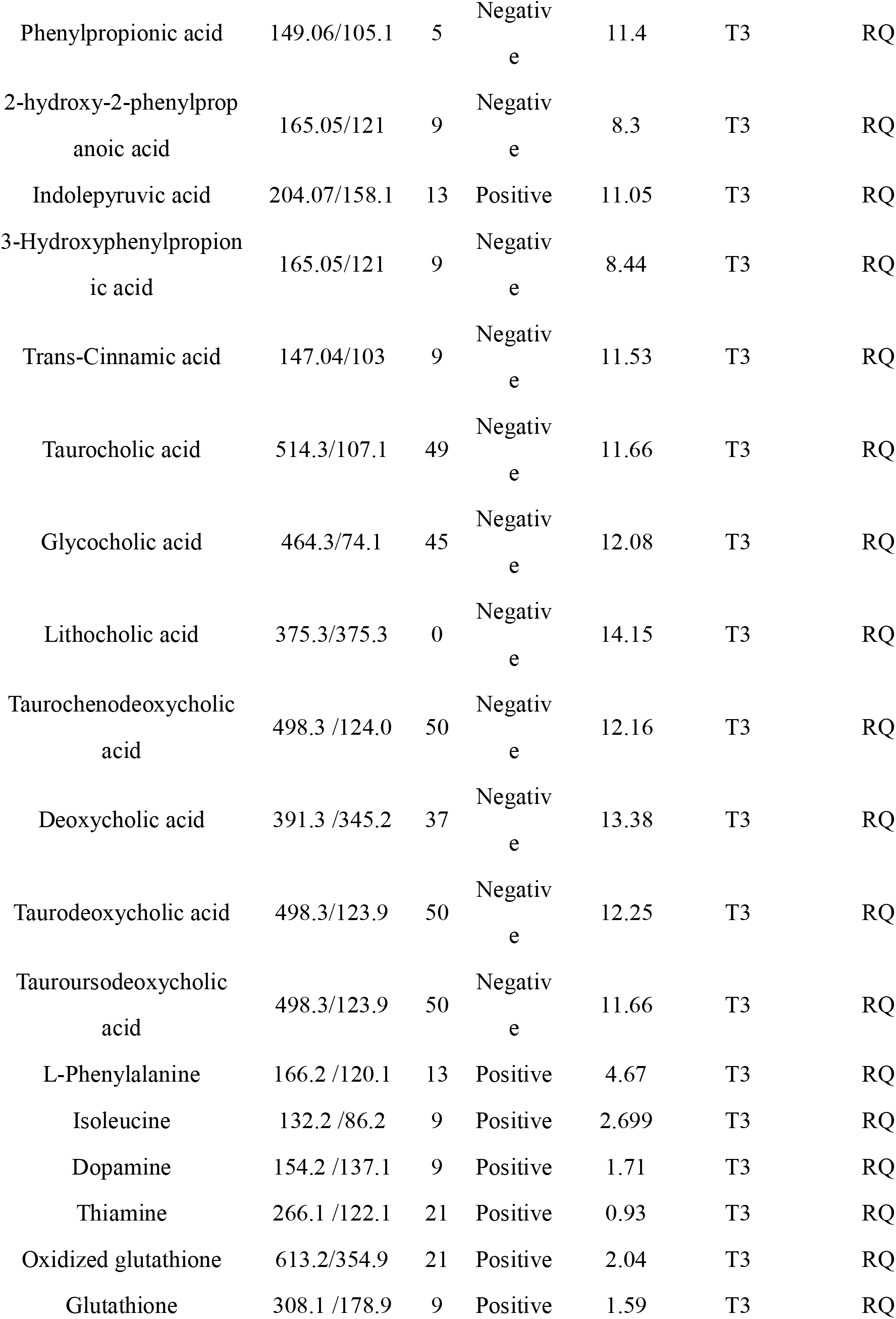
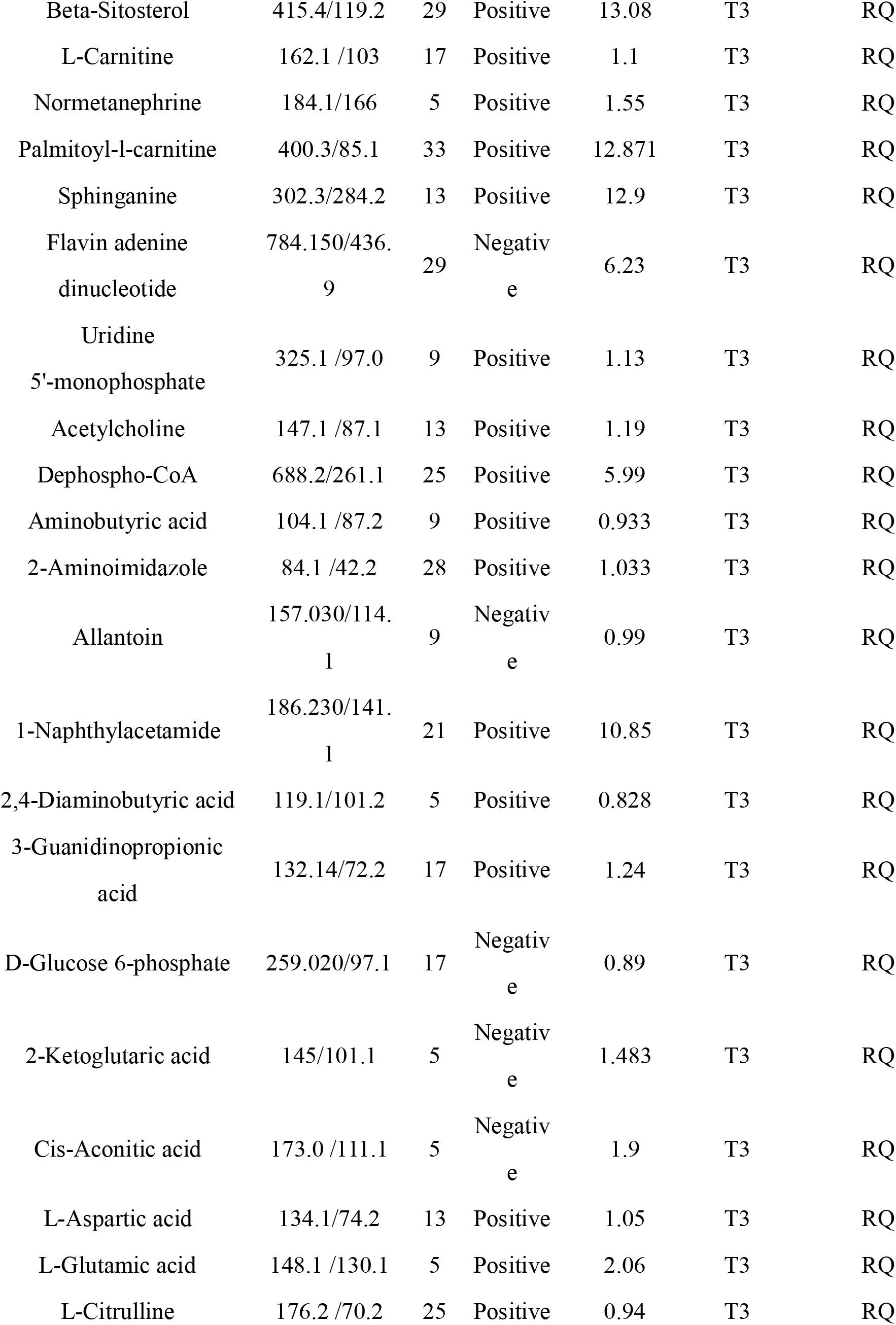
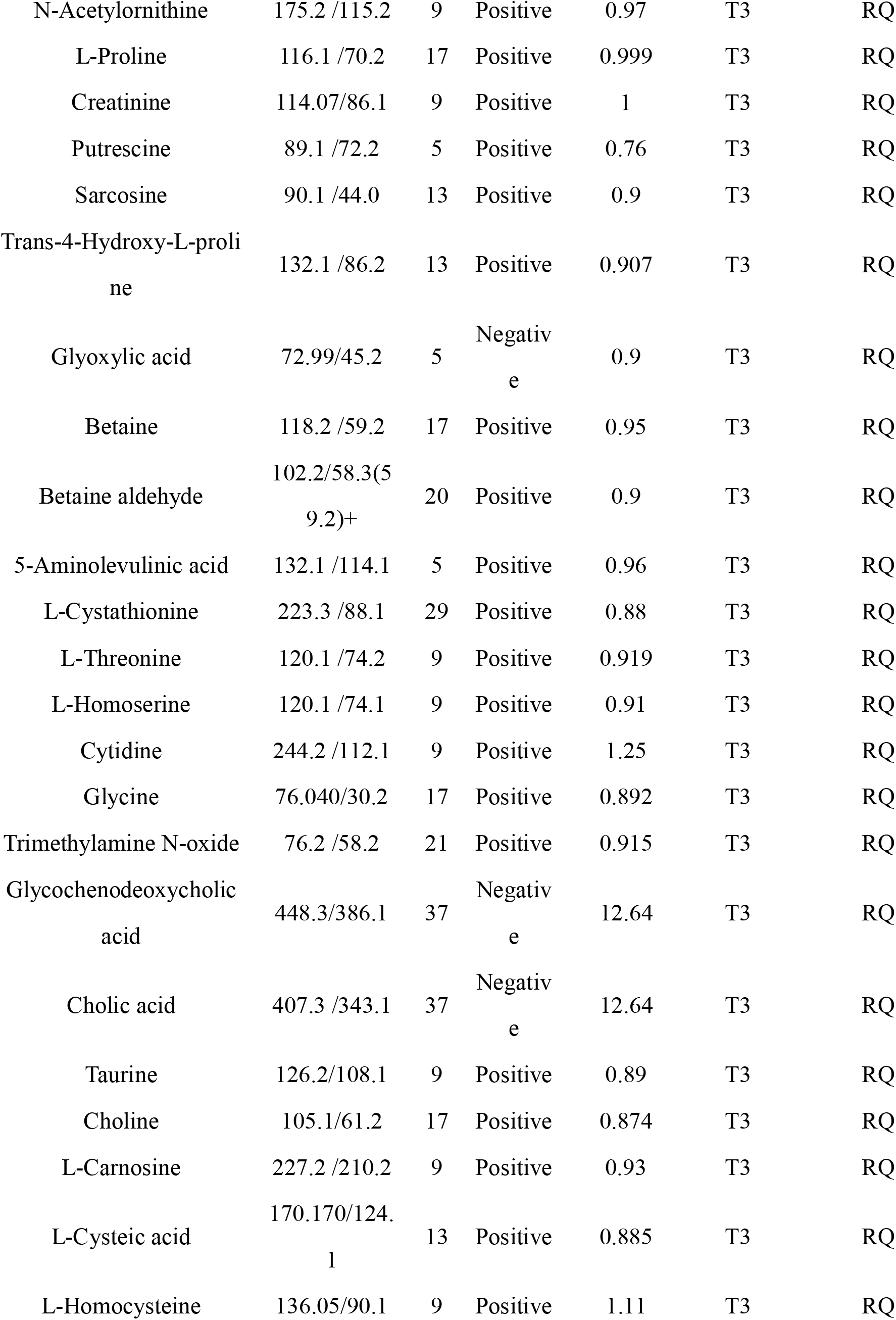
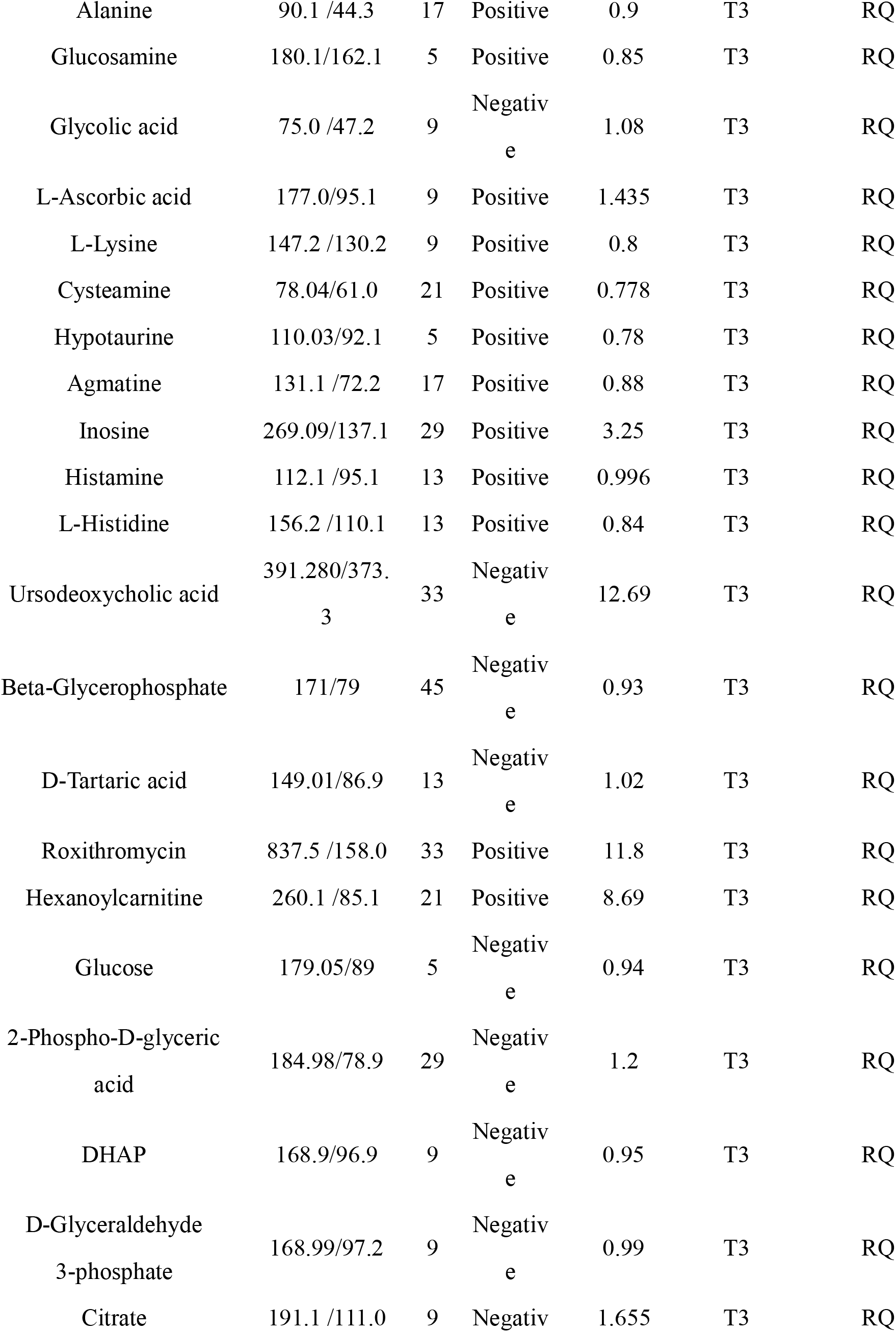
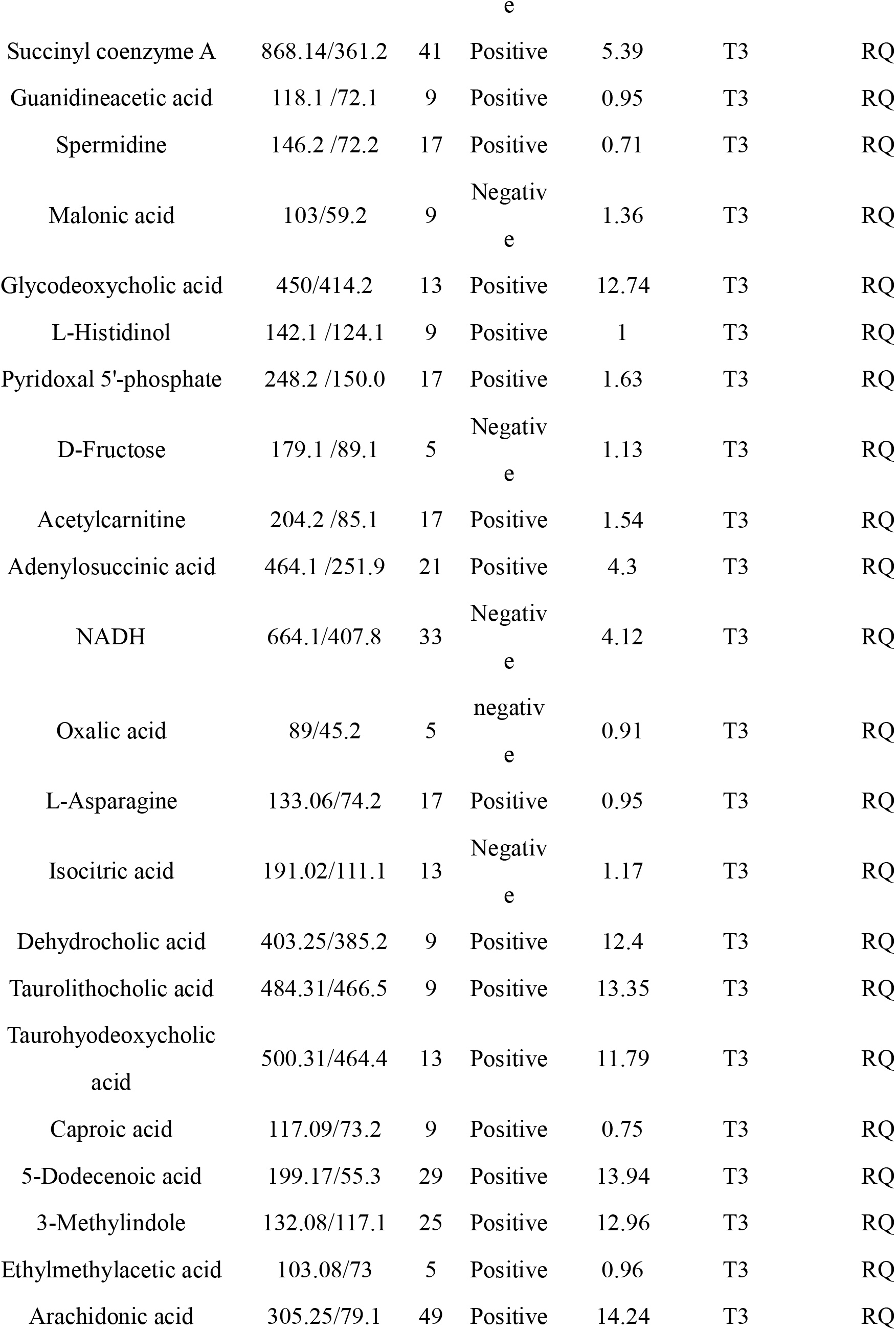
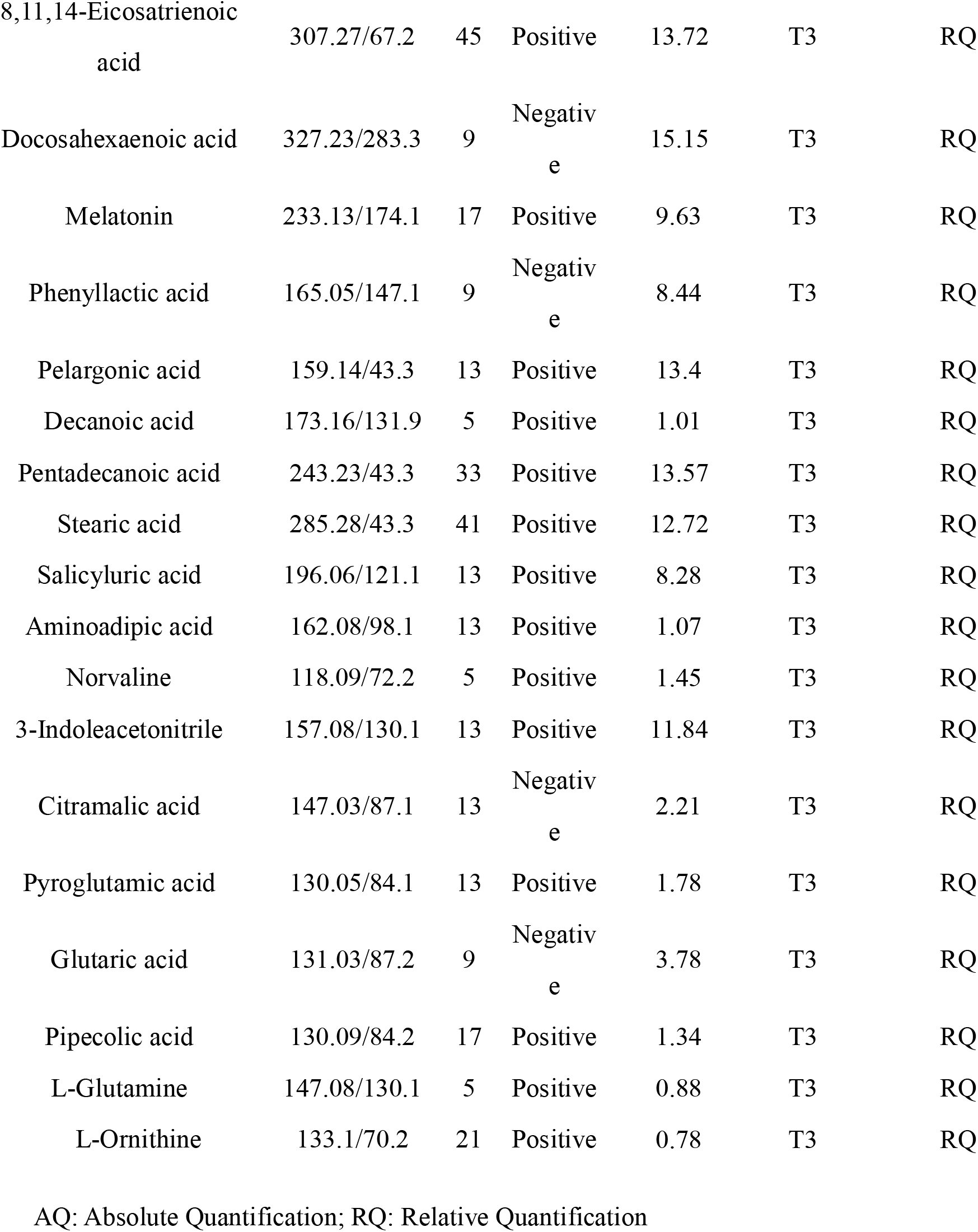
LC-TQ dynamic-MRM parameters of metabolomes

**Table S2.**
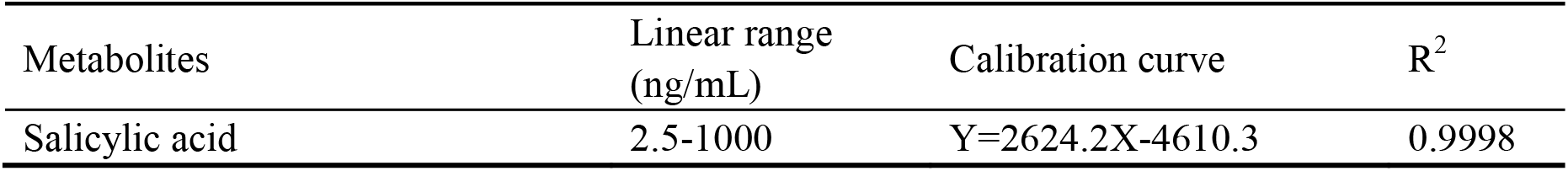
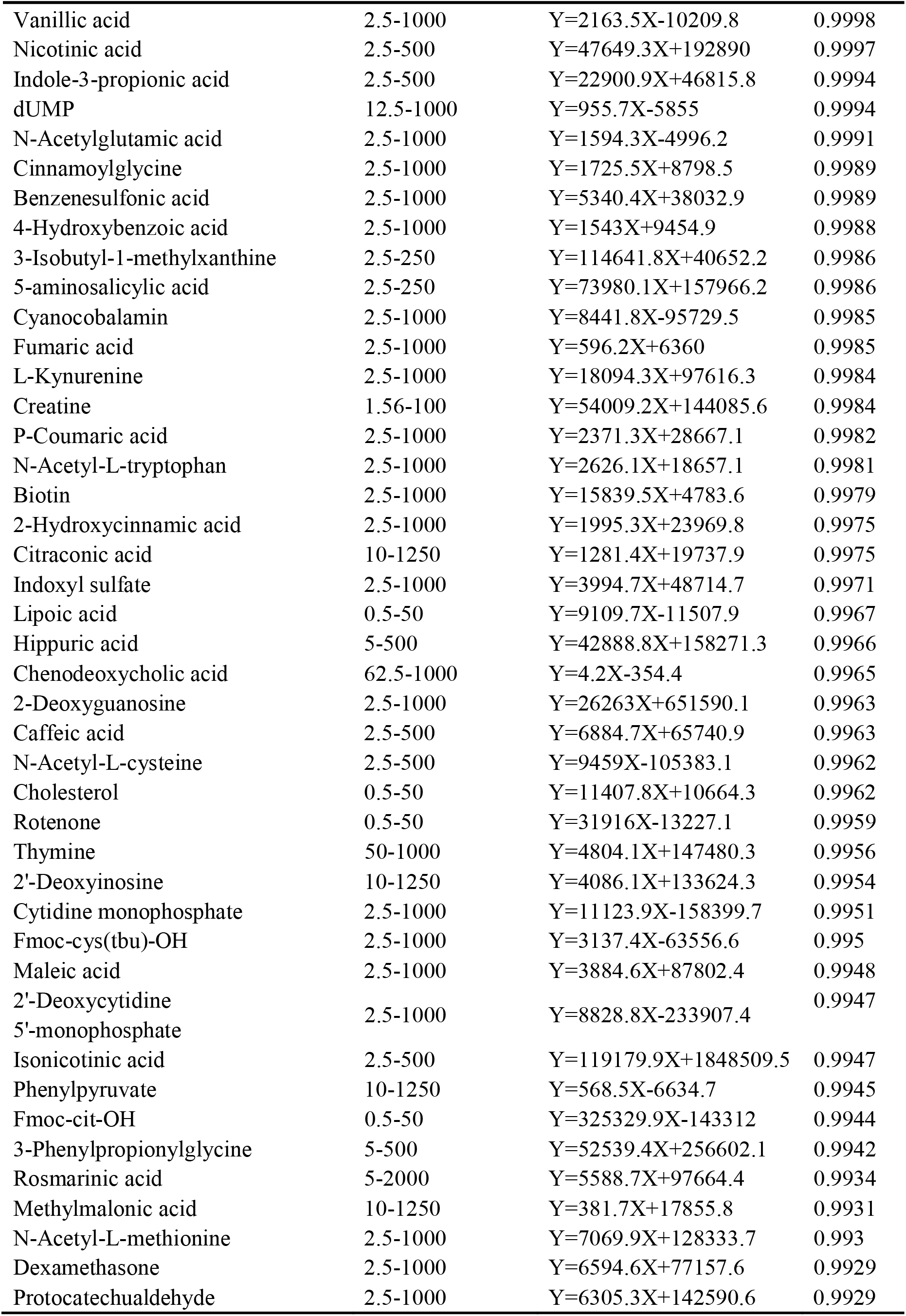
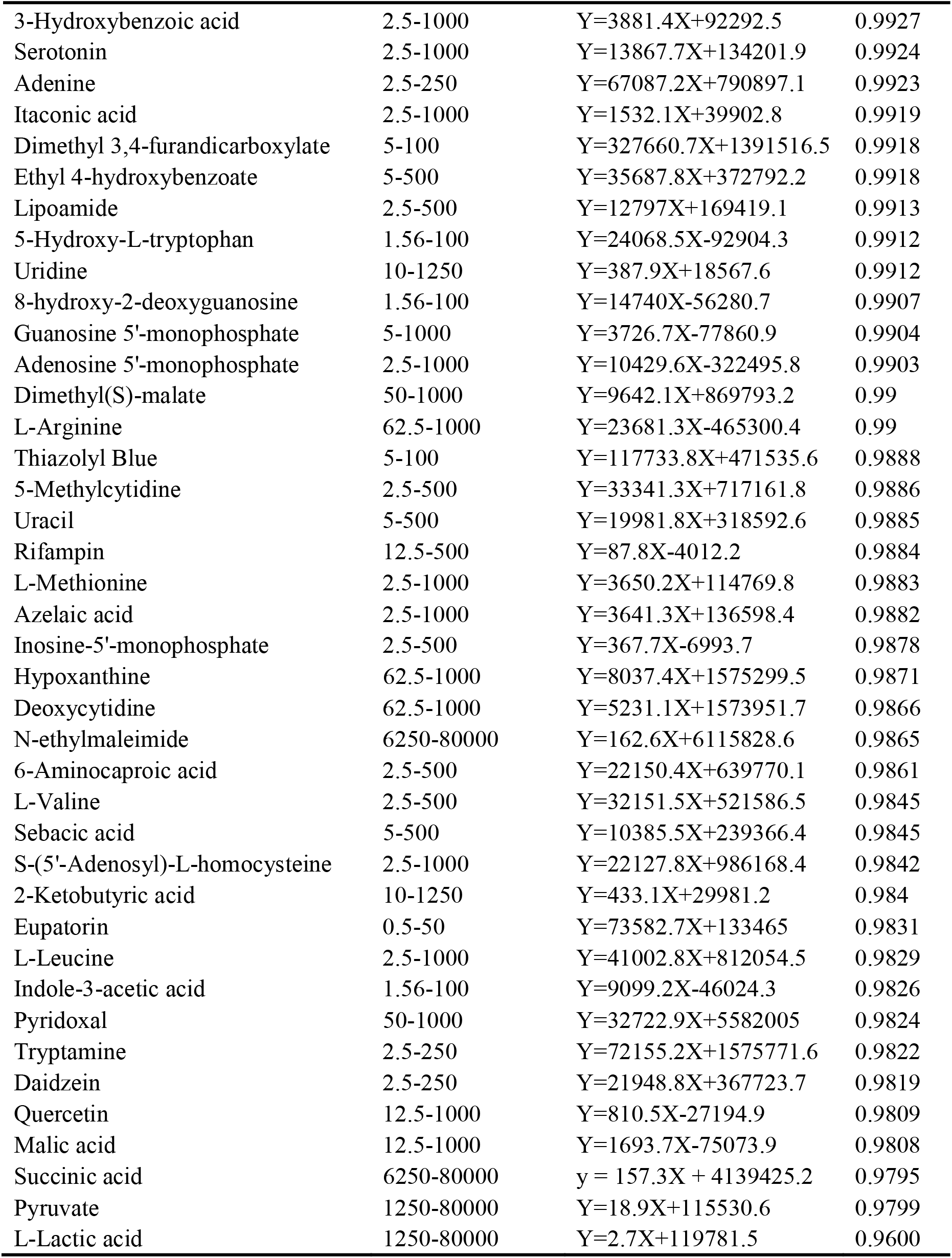
Absolute determination of 84 key metabolites with calibration-curve

**Table S3.**
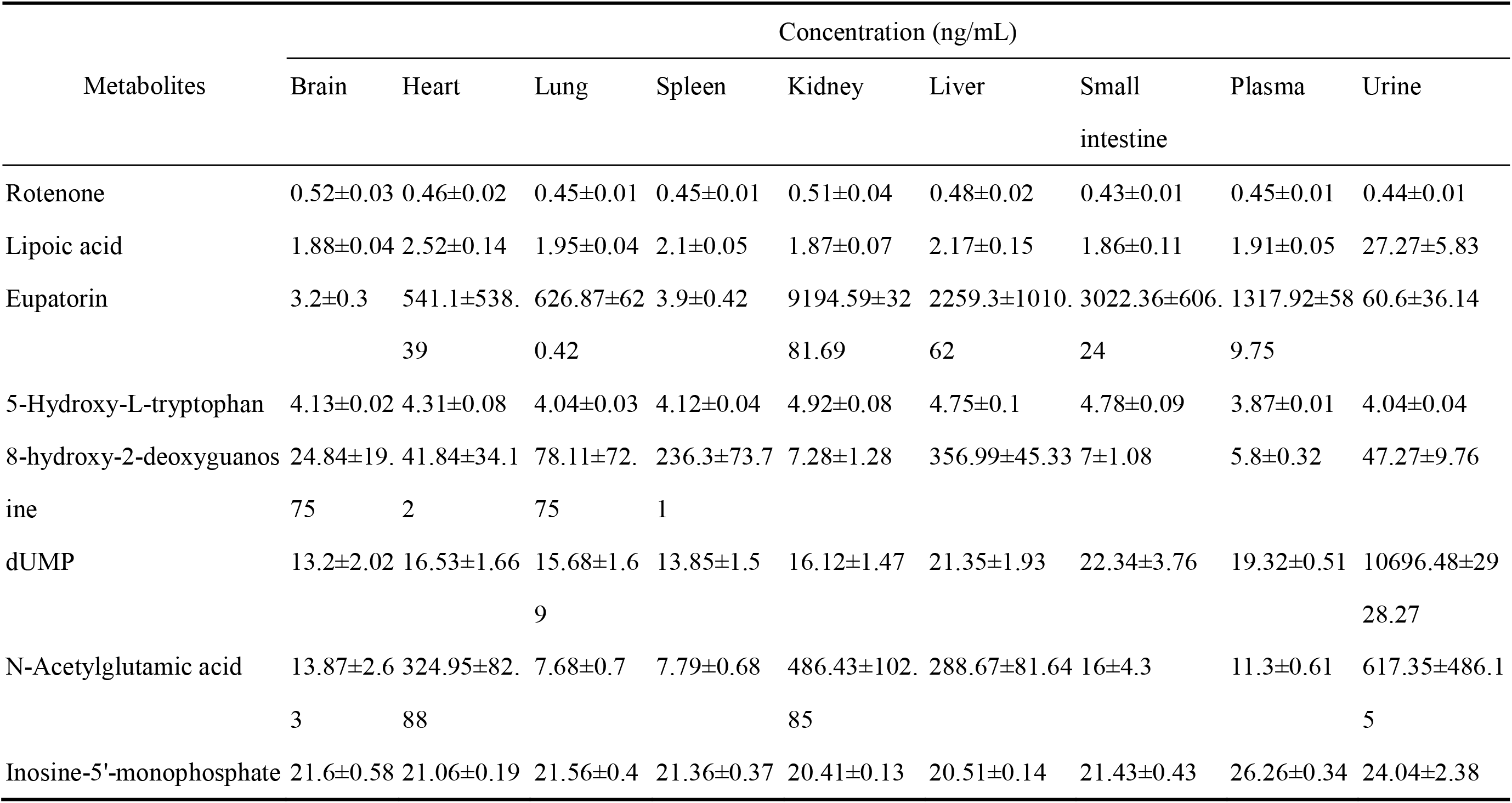
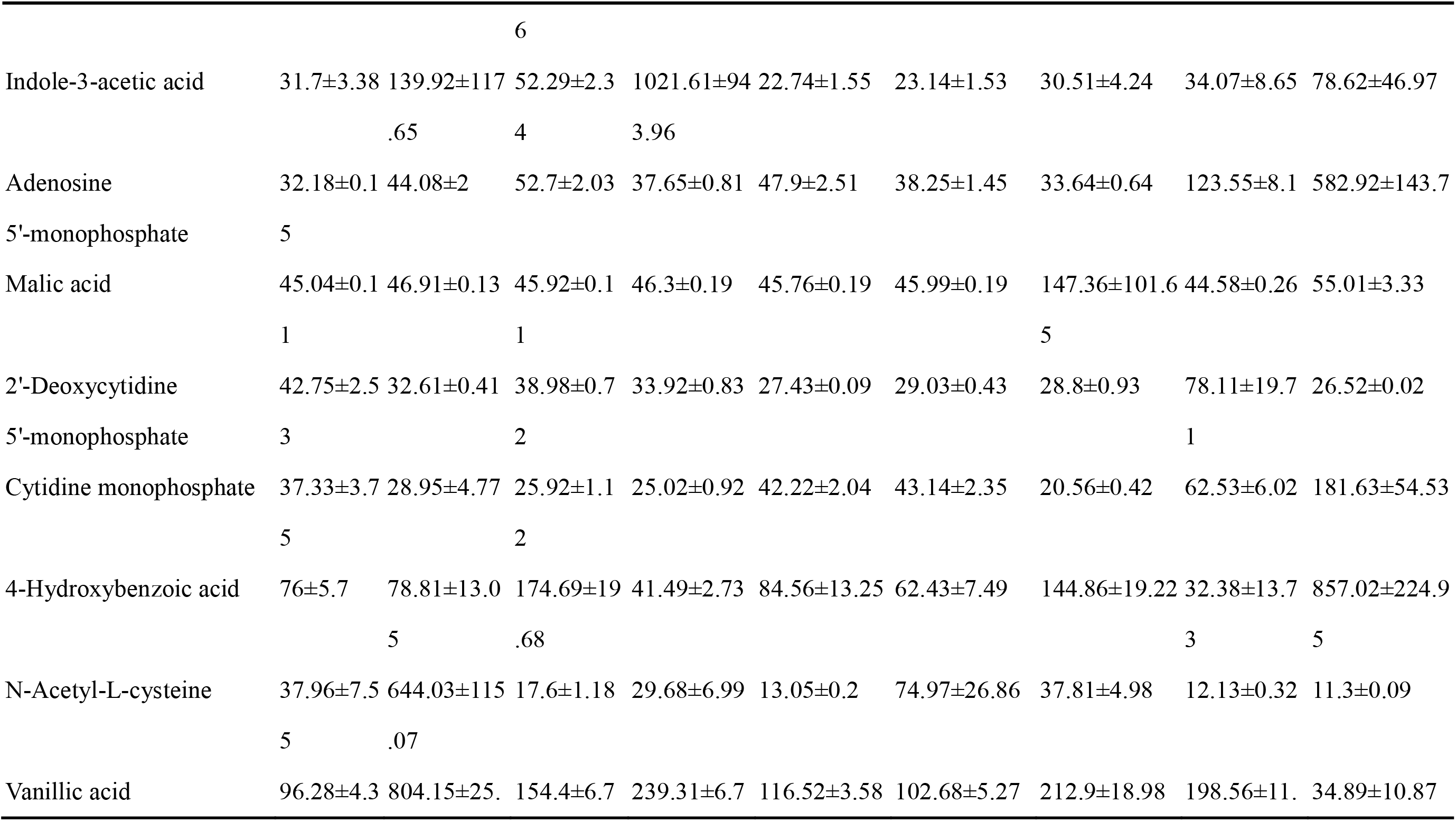
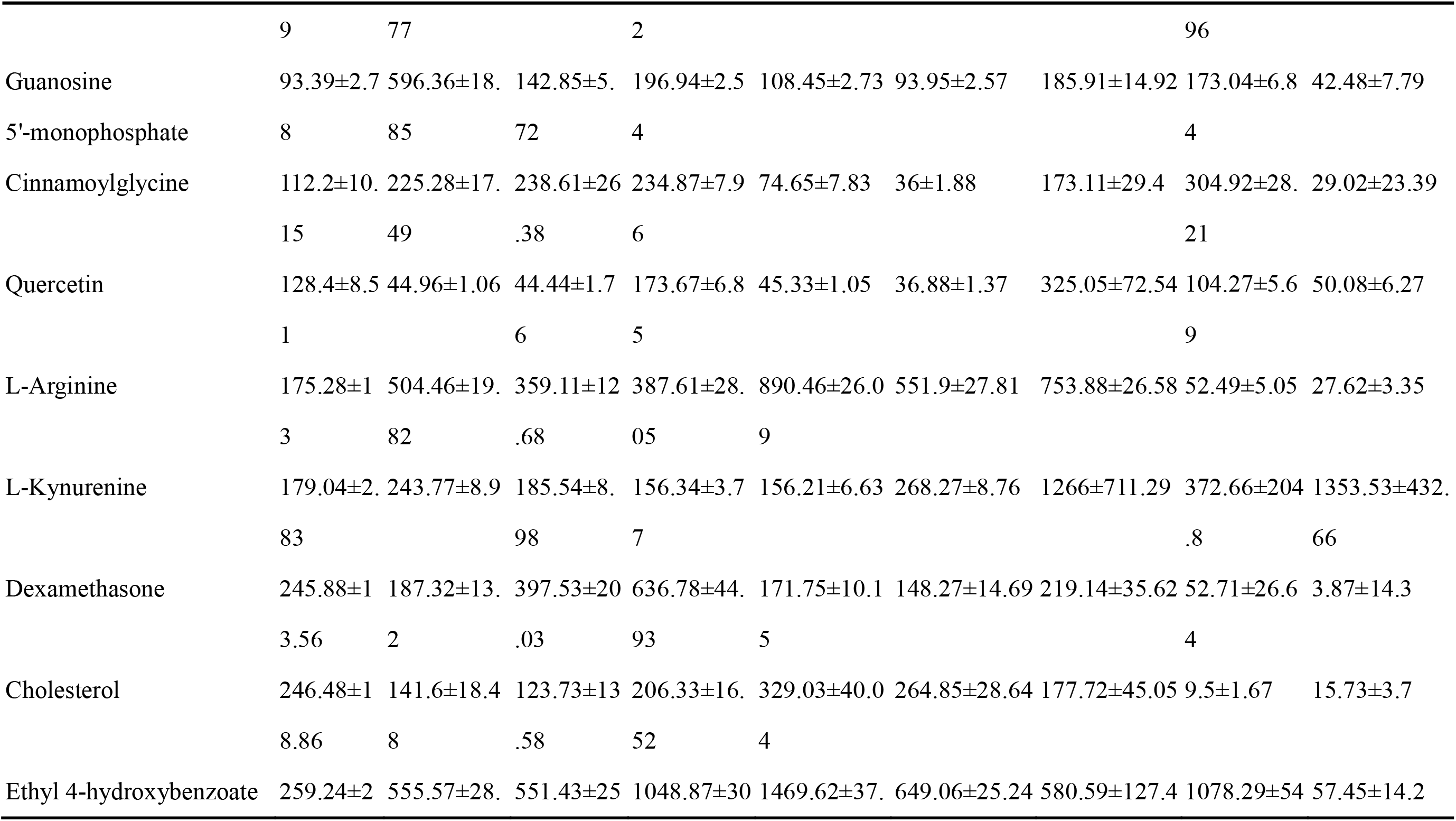
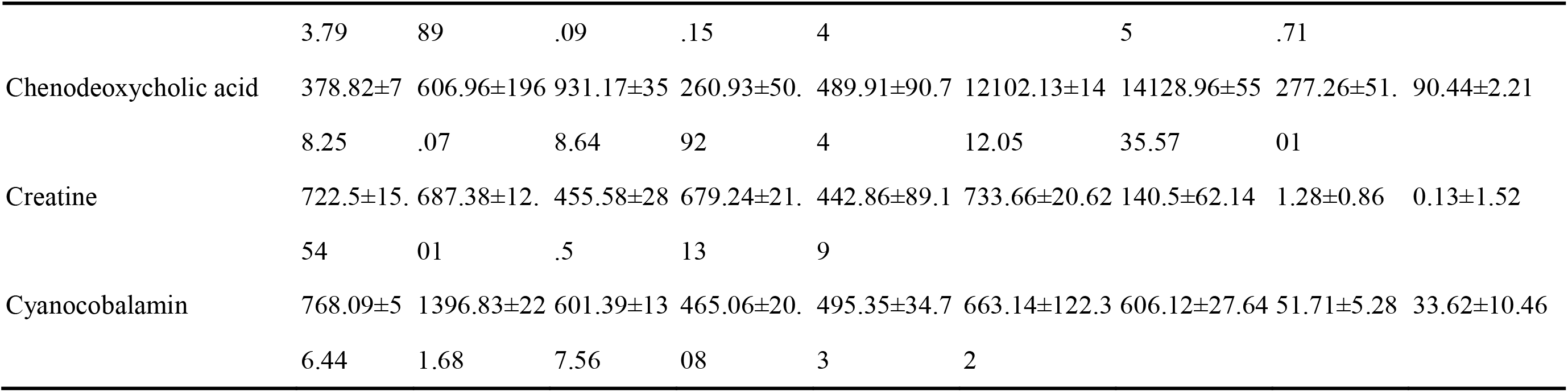
Concentration of representative 27 key metabolites in plasma, urine and a diverse tissues of rats (n=6)

## References

1 Jang, C., Chen, L. & Rabinowitz, J. D. Metabolomics and Isotope Tracing. Cell 173, 822–837, doi:10.1016/j.cell.2018.03.055 (2018).

2 Boulange, C. L., Neves, A. L., Chilloux, J., Nicholson, J. K. & Dumas, M. E. Impact of the gut microbiota on inflammation, obesity, and metabolic disease. Genome Med 8, 42, doi:10.1186/s13073-016-0303-2 (2016).

3 Geiger, R. et al. L-Arginine Modulates T Cell Metabolism and Enhances Survival and Anti-tumor Activity. Cell 167, 829–842.e813, doi:10.1016/j.cell.2016.09.031 (2016).

4 Oliver, S. G., Winson, M. K., Kell, D. B. & Baganz, F. Systematic functional analysis of the yeast genome. Trends Biotechnol 16, 373–378, doi:Doi 10.1016/S0167-7799(98)01214-1 (1998).

5 Nicholson, J. K., Lindon, J. C. & Holmes, E. ‘Metabonomics’: understanding the metabolic responses of living systems to pathophysiological stimuli via multivariate statistical analysis of biological NMR spectroscopic data. Xenobiotica 29, 1181–1189, doi:Doi 10.1080/004982599238047 (1999).

6 Winder, C. L., Dunn, W. B. & Goodacre, R. TARDIS-based microbial metabolomics: time and relative differences in systems. Trends Microbiol 19, 315–322, doi:10.1016/j.tim.2011.05.004 (2011).

7 Want, E. J. et al. Global metabolic profiling of animal and human tissues via UPLC-MS. Nature protocols 8, 17–32, doi:10.1038/nprot.2012.135 (2013).

8 Want, E. J. et al. Global metabolic profiling procedures for urine using UPLC-MS. Nature protocols 5, 1005–1018, doi:10.1038/nprot.2010.50 (2010).

9 Kuo, M. T., Savaraj, N. & Feun, L. G. Targeted cellular metabolism for cancer chemotherapy with recombinant arginine-degrading enzymes. Oncotarget 1, 246–251, doi:10.18632/oncotarget.135 (2010).

10 Luo, X. et al. Mass spectrometry and associated technologies delineate the advantageously biomedical capacity of siderophores in different pathogenic contexts. Mass Spectrometry Reviews 0, doi:doi:10.1002/mas.21577.

11 Cai, Y., Weng, K., Guo, Y., Peng, J. & Zhu, Z.-J. An integrated targeted metabolomic platform for high-throughput metabolite profiling and automated data processing. Metabolomics 11, 1575–1586, doi:10.1007/s11306-015-0809-4 (2015).

12 Wei, R., Li, G. D. & Seymour, A. B. High-Throughput and Multiplexed LC/MS/MRM Method for Targeted Metabolomics. Analytical chemistry 82, 5527–5533, doi:10.1021/ac100331b (2010).

13 Lv, H., Palacios, G., Hartil, K. & Kurland, I. J. Advantages of tandem LC-MS for the rapid assessment of tissue-specific metabolic complexity using a pentafluorophenylpropyl stationary phase. Journal of proteome research 10, 2104–2112, doi:10.1021/pr1011119 (2011).

14 Su, Q., Guan, T., He, Y. & Lv, H. Siderophore Biosynthesis Governs the Virulence of Uropathogenic Escherichia coli by Coordinately Modulating the Differential Metabolism. Journal of proteome research 15, 1323–1332, doi:10.1021/acs.jproteome.6b00061 (2016).

15 Lv, H., Hung, C. S., Chaturvedi, K. S., Hooton, T. M. & Henderson, J. P. Development of an integrated metabolomic profiling approach for infectious diseases research. Analyst 136, 4752–4763, doi:10.1039/c1an15590c (2011).

16 Lv, H. et al. Ingenuity pathways analysis of urine metabonomics phenotypes toxicity of gentamicin in multiple organs. Mol Biosyst 6, 2056–2067, doi:10.1039/c0mb00064g (2010).

17 Domingo-Almenara, X. et al. XCMS-MRM and METLIN-MRM: a cloud library and public resource for targeted analysis of small molecules. Nat Methods 15, 681–684, doi:10.1038/s41592-018-0110-3 (2018).

18 Rao, Z. et al. Development of a dynamic multiple reaction monitoring method for determination of digoxin and six active components of Ginkgo biloba leaf extract in rat plasma. Journal of chromatography. B, Analytical technologies in the biomedical and life sciences 959, 27–35, doi:10.1016/j.jchromb.2014.03.028 (2014).

